# Meningococcal vaccine Bexsero elicits a robust cellular immune response that targets but is not consistently protective against *Neisseria gonorrhoeae* during murine vaginal infection

**DOI:** 10.1101/2024.09.08.611931

**Authors:** Joseph J. Zeppa, Jamie E. Fegan, Pauline Maiello, Epshita A. Islam, Isaac S. Lee, Christine Pham, Laura-Lee Caruso, Scott D. Gray-Owen

**Affiliations:** Department of Molecular Genetics, Temerty Faculty of Medicine, University of Toronto, Toronto, ON, Canada; Department of Microbiology and Molecular Genetics, University of Pittsburgh School of Medicine, Pittsburgh, PA, USA; Center for Vaccine Research, University of Pittsburgh School of Medicine, Pittsburgh, PA, USA; Department of Biochemistry, Temerty Faculty of Medicine, University of Toronto, Toronto, ON, Canada

## Abstract

Retrospective epidemiological studies suggest that the licensed serogroup B meningococcal vaccine 4CMenB (Bexsero) provides some protection against the closely related pathogen *Neisseria gonorrhoeae* in humans. This result has been replicated in murine models of gonococcal colonization, with a gonococci-reactive humoral response and more rapid clearance of vaginal infection. However, immunization with Bexsero consistently elicits a robust humoral response but does not protect all individuals, so the correlates of protection remain undefined. Herein, we exploit the fact that Bexsero promotes clearance in only a subset of immunized mice to perform a broad analysis of the adaptive response in animals that are or are not protected. We observe that Bexsero vaccination induces high levels of anti-neisserial antibodies in both serum and the vaginal lumen, and a robust cellular response highlighted by an increase in both conventional naïve and memory populations as well as unconventional lymphocyte subsets. Multiplex and flow cytometry results show that Bexsero vaccination generates a robust, multi-faceted cytokine response that spans numerous T cell subsets (T_H_1, T_H_2, T_reg_ and T_H_17 responses) and that non-T non-B lymphocytes play an important role in this response, as indicated by an unbiased principal component analysis. Together, this work provides the first comprehensive analysis of the robust humoral and complex cellular response to Bexsero so as to reveal the effector mechanisms that may contribute to immunity against vaginal gonococcal infection.

## Introduction

*Neisseria gonorrhoeae* (Ngo), which causes the sexually transmitted infection gonorrhea, is an ongoing burden globally and an important bacterial pathogen responsible for over 82 million new infections in 2020^1^. Although still treatable with antibiotics, the emergence and rapid evolution of antimicrobial resistant strains is raising the possibility of untreatable gonorrhea^2^. Uncomplicated urogenital infection by Ngo does not typically lead to the generation of a protective immune response, allowing for repeated infections^3^. Due to this absence of immunity, it remains unknown as to what immune responses can confer protection and thus it remains unclear what type of correlates can be used to guide the development of a new gonococcal vaccine^4^.

The four component meningococcal vaccine 4CMenB (marketed as Bexsero by GSK), is a licensed vaccine that was developed to protect against serogroup B strains of the pathogen *Neisseria meningitidis* (Nme)^5^. This multicomponent vaccine is comprised of outer membrane vesicles (OMVs) from Nme strain NZ98/254 mixed with recombinant meningococcal heparin binding antigen (NHBA), neisserial adhesin A (NadA), and factor H binding protein (fHbp), along with two proteins fused to NHBA and fHbp (GNA1030 and GNA2091, respectively) to increase bactericidal titres^6^. Serological analysis of individuals who have been immunized with this vaccine has identified antibodies that cross react with Ngo antigens that are conserved between these organisms^7,8^, including OMV-derived antigens and the recombinant NHBA, GNA2091 and GNA1030, although the latter two are not believed to be surface exposed in Ngo^7^. Even more remarkable is that several epidemiological studies have found that individuals vaccinated with Bexsero have a reduced rate of infection by Ngo^9–11^. Vaccine efficacy is estimated to range between 33 – 44% in humans, a finding that has recently prompted some governments to recommend vaccinating individuals at high risk of contracting gonorrhea with Bexsero^12^.

Given the remarkable finding that Bexsero confers some protection against Ngo infection in humans, it is satisfying that this vaccine also confers protection against gonococcal infection in the well-established mouse vaginal infection model. In this case, Bexsero accelerated gonococcal clearance from the lower genital tract and elicited cross-reactive antibodies to some of same gonococcal components recognized by Bexsero-immune human serum^13^, establishing the relevance of this model for further investigation of vaccine-induced mechanisms of protection. In this study, we have taken a comprehensive approach to analyze the cellular and humoral immune responses for each individual animal to identify potential immune correlates of protection. ELISAs, Western blots, and serum bactericidal assays were performed to compare the humoral response against Ngo in control versus vaccinated animals that were or were not protected; flow cytometry was performed on spleen and genital tract lymphocytes to assess immune populations and cellular cytokine responses; and multiplex assays were performed to assess cytokines and chemokines from vaccine-stimulated lymphocytes from the vaginal lumen, genital tract tissues and spleen prior to and during gonococcal infection. Finally, principal component analyses were performed on all data sets to determine what factors were driving the variability in our data and determine if immune correlates of protection could be revealed by this study. This unbiased approach demonstrates that the vaccine elicits consistently high humoral response that does not explain the different level of protection in these animals and reveals effector mechanisms that may combine to contribute to immunity against vaginal infection by Ngo.

## Results

### Bexsero reduces the duration of colonization and bacterial load of *Neisseria gonorrhoeae* in a murine vaginal infection model

To establish that the previously reported protection that Bexsero conferred against mouse vaginal colonization by Ngo could be replicated in our study, we performed a preliminary infection experiment by vaginal inoculation of female BALB/c mice to monitor protection and allow us to power this study. Consistent with the previous work, we saw a significant reduction in the percentage of colonized mice that were Bexsero vaccinated compared to the control group. Based upon this, we powered our study using percent colonization data from day six because it was the first day that we observed equivocal protection data between our Ngo infection model and that observed in human retrospective studies (approximate vaccine efficacy of 40%) and we wanted to identify early effectors of protection rather than monitor immunity post-infection. Using a Type I error rate of 0.05, 80% power, and an enrollment rate of 1:1 it was determined that we would need a minimum sample size of 14 animals per group to detect a statistical difference in colonization. Additionally, this balance of animals would also give us the opportunity to perform complex cellular analyses to search for potential immune correlates of protection as a secondary analysis.

The mice were randomized 1:1 into either alum control or Bexsero group, vaccinated at day 0, 21 and 42, and then staged to allow infection ∼1 × 10^7^ colony forming units (CFU) of Ngo strain FA1090 during estrus. Vaginal lavages were collected daily to assess colonization and enumerate bacterial burden (Figure 1a). As in our pilot study, a statistically significant reduction in the percentage of animals colonized (p = 0.0284, Figure 1b). Comparison of vaginal bacterial load between Bexsero and alum on each day also revealed a significant reduction in Bexsero vaccinated animals on days three and four (p = 0.0079 and 0.0041, respectively; Figure 1c). We also observed a statistically significant reduction in the number of CFU positive days in our Bexsero group (p = 0.0277, Figure 1d), however, the cumulative CFU was not statistically different (p = 0.0825, Figure 1e) but was trending lower in the Bexsero group. Within the Bexsero group it was noted that there were approximately 44% (8/18) of animals that had no bacterial recovery after day three post infection; we labelled these as ‘protected’ animals. When this group was separated from the ‘not protected’ Bexsero immunized animals and compared to the alum control group, it was noted that the ‘protected’ group had statistically fewer days colonized and lower cumulative CFU, whereas the unprotected animals were not statistically different from controls (Supplemental Figures 1a and b). We also noted a significant correlation (p = 0.0001; R^2^ = 0.3971) between both the number of colonized days and cumulative CFU between the protected and all other animals in the study (Supplemental Figure 1c).

**Figure 1.**
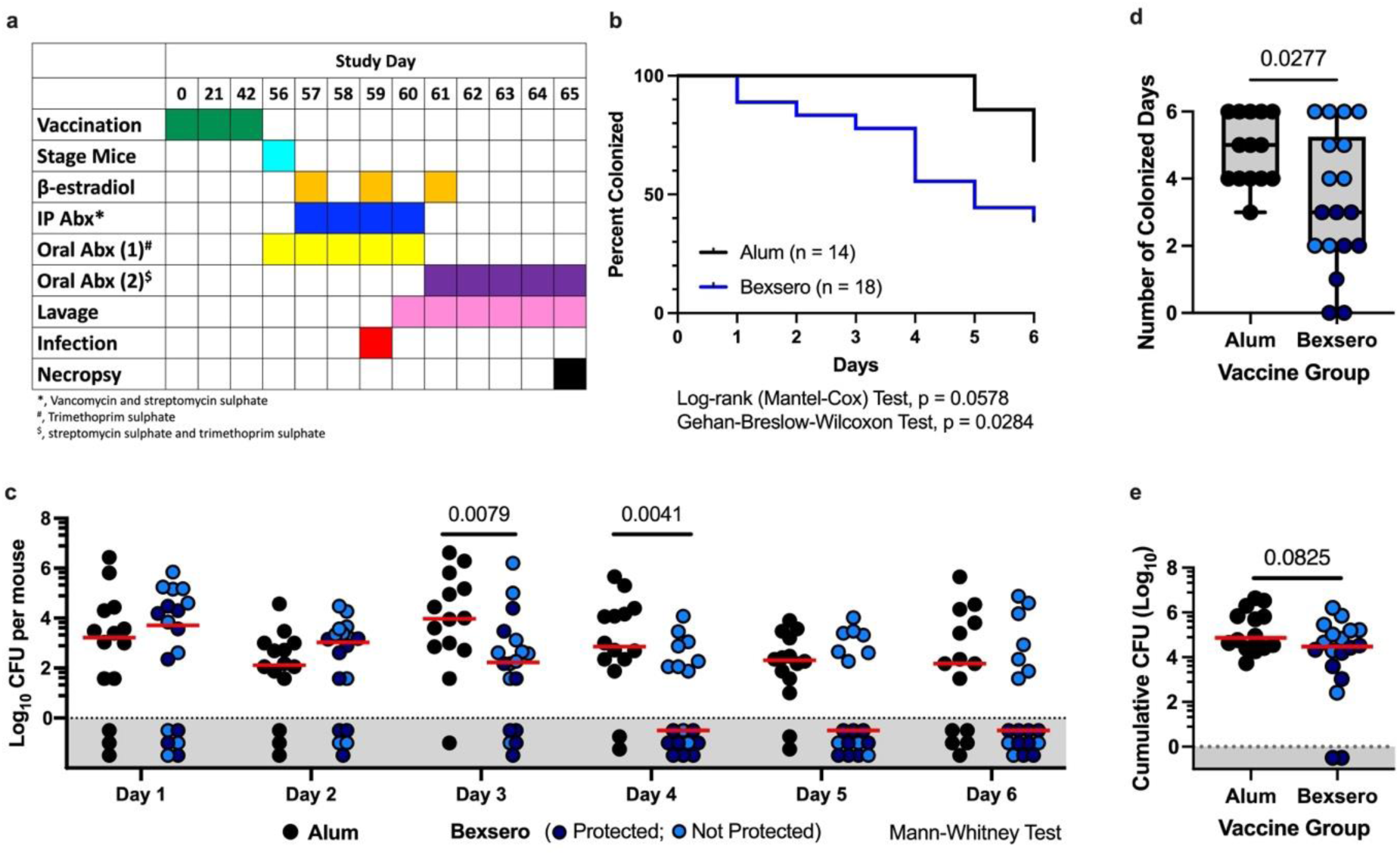
Bexsero vaccination protects mice from vaginal gonococcal infection, speeds up clearance and reduces bacterial load. (**a**) The vaccination and challenge study outline. (**b**) Kaplan-Meier curve showing the percentage of animals colonized in each group (Bexsero, blue line; alum control, black) on each day post infection. P values for Log-rank (Mantel-Cox) and Gehan-Breslow-Wilcoxon statistical tests are shown. (**c**) The vaginal bacterial load recovered from each animal on each day. (**d**) The total number of days colonized for each animal. The Box plot denotes the median and upper/lower quartiles within the grey area with the whiskers extending to the minimum and maximum values. (**e**) The cumulative bacterial load recovered from each animal throughout the experiment. Each symbol represents one animal. The alum control group is shown in black. The Bexsero-vaccinated group is either blue (**b**) or separated into dark blue circles (Bexsero-vaccinated protected; no bacterial recovered from vaginal lavages on at minimum days four through six) or light blue circles (not protected Bexsero-vaccinated animals; bacteria recovered from vaginal lavage on one or more of the last three days of the experiment) (**c** – **e**). Red horizontal lines denote the median and symbols within the grey area denote no Ngo recovery (**c** and **e**). Mann-Whitney statistical test performed, and p values listed (**c** – **e**).

### Bexsero vaccination elicited significant anti-neisserial antibody titers with notable mouse to mouse variability in the bacterial components recognized

To compare the humoral response between our two groups, ELISAs were performed using pre-infection (post-vaccination) serum and vaginal lavage against whole cell heat-inactivated Nme NZ98/254 (the parental strain of the OMVs in Bexsero) and Ngo FA1090 (challenge strain used in this study). As expected, we observed a significantly higher serum IgG titer against both bacterial strains (p = 0.0001) in our Bexsero-vaccinated animals compared to control (Supplemental Figure 2a and b), though titres against the Ngo challenge strain were approximately 20 times lower than they were against Nme (median values of 61.29 μg mL^-1^ and 1,238.93 μg mL^-1^, respectively). For vaginal IgA, titers against the meningococcal strain were significantly higher in Bexsero- immunized animals (p = 0.0289) but titers against the challenge gonococcal strain were not (Supplemental Figure 2c and d). Additionally, there was no difference in titers observed between our protected and not protected Bexsero animals (Supplemental Figure 2). We also noted that there was no correlation with total serum IgG or mucosal IgA antibody titers and the cumulative CFU or number or colonized days (Supplemental Figure 3).

To further categorize the IgG response, IgG isotypes in terminal serum against inactivated Nme NZ98/254 and Ngo FA1090 were quantified. Significant binding of all four isotypes (IgG1, IgG2a, IgG2b, and IgG3) was detected for the Bexsero group against Nme NZ98/254 (Supplemental Figure 4a–d, p<0.0001) and significant, albeit lower, IgG1, IgG2a, and IgG2b were detected against Ngo FA1090 (Supplemental Figure 4e–h, p<0.0001). Notably, IgG2a and IgG2b were the predominant classes bound to Nme NZ98/254, while cross-reactive antibodies against Ngo FA1090 were mainly IgG1. A negative correlation between IgG2b signal against Nme NZ98/254 was observed for both number of colonized days (p = 0.0332) and cumulative CFU (P = 0.0368). However, IgG2b signal against the challenge strain Ngo FA1090 itself did not show a significant interaction and there was also no difference for any of the Isotypes in protected versus not protected Bexsero animals (Supplemental Figure 4).

Considering that Bexsero is a multicomponent vaccine, we performed Western blots with terminal immune (post immunization and challenge) serum against whole bacterial lysates of both Nme NZ98/254 and Ngo FA1090 to examine if animal-to-animal variability exists in which bacterial components are recognized. Several bands that resolved between 25 to 125 kDa were recognized, with noticeable inter-animal variability in both intensity and number of bands (Supplemental Figure 5a and b). Intensities of each distinct band against Ngo FA1090 at two exposures were measured and analyzed against number of colonized days or cumulative CFU for potential correlations (Supplemental Figure 5c). A moderate negative correlation with number of colonized days was observed for a ∼85 kDa (p = 0.0220) and a ∼65 kDa band (p = 0.0196); however, there was no significant difference in intensities between protected versus not protected animals (Supplemental Figure 5d and e).

Finally, serum bactericidal activity (SBA) assays were performed against Ngo FA1090 using heat inactivated 2-fold serially diluted terminal serum and either 2.5% baby rabbit serum (Supplemental Figure 6a and b) or 15% normal human serum (Supplemental Figure 6c and d) as the complement source. Only 5 out of 32 samples produced an SBA titre (dilution at which 50% killing occurs relative to no antibody control) with human serum, to which FA1090 is highly resistant^14^, while 29 out of 32 samples produced an SBA titre using baby rabbit serum. In both assays, there was no significant difference in CFU recovered at each dilution or in SBA titre among adjuvant control and Bexsero-immunized groups (Supplemental Figure 6).

Taken together, these studies suggest high levels of animal-to-animal variability in the humoral response elicited by Bexsero. While none of our humoral readouts correlated with protection status, a few promising correlations with number of days colonized were noted (Western blot bands 3 and 6 [Supplemental Figure 5d and e]).

### T cells and CD4+ memory cells increase in Bexsero-vaccinated mice

On day six post-infection, mice were euthanized, necropsied, and immune cells were isolated from both the spleen and female genital tract (FGT) and subjected to flow cytometric analysis to determine if Bexsero vaccination had an impact on specific immune cell subsets, and particularly whether the protected and unprotected groups differed. B cells (CD19+), T cells (CD3+), T cells subsets (CD4+ and CD8α+), and non-T non-B lymphocytes (CD3- CD19-) were assessed (Figure 2a and b). We noticed a statistically significant increase in CD4+ T cell frequencies in the spleen in our Bexsero vaccinated group (p = 0.0109), with subsequent decreases in CD8α+ (p = 0.0092) and CD19+ cells (p = 0.0012; Figure 2c). In the genital tract, we noted an increase in overall CD3+ T cell frequences in the Bexsero group (p = 0.0092), with a decrease in the CD8α+ T cells (p = 0.0215; Figure 2d). There were no differences in total cell numbers per gram of tissue in the spleen (Figure 2e), whereas there were statistically more CD3+ T cells (p = 0.0384) in the FGT (Figure 2f); this difference did not differentiate between protected and unprotected animals.

**Figure 2.**
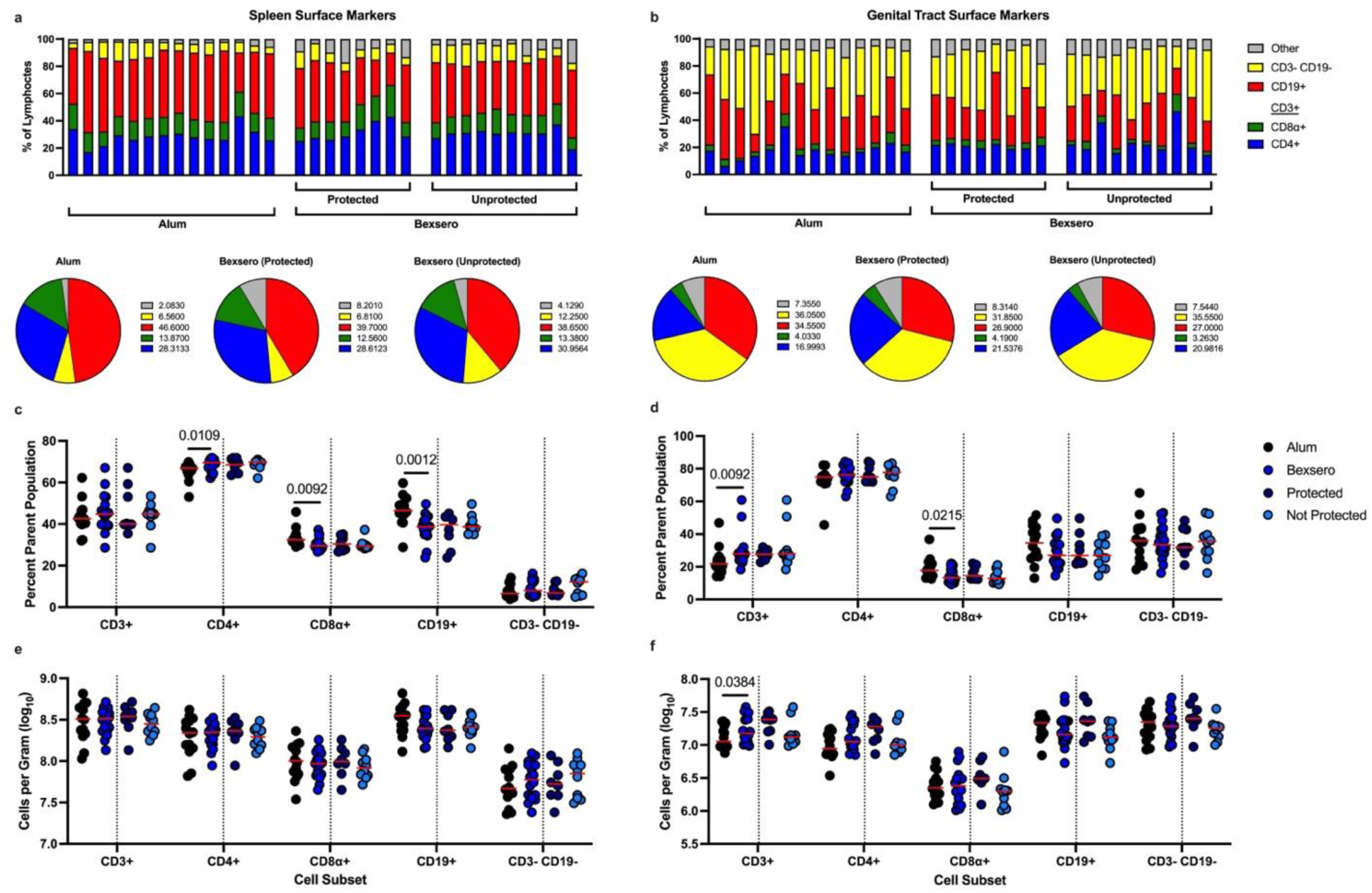
Splenic CD4+ T cell frequences and genital tract CD3+ T cells increase after immunization with Bexsero. After three vaccinations of Bexsero or alum and six days post Ngo vaginal infection female BALB/c mice were euthanized and necropsied, and frequencies plus total cell counts per gram of tissue were assessed in the spleen and the genital tract using flow cytometry. The frequency of indicated cell subsets are shown in the spleen (**a**, Top) or the genital tract (**b**, Top), with animals divided into their vaccination group and Bexsero-protection status. Each bar represents one animal. Pie charts denote the median values of each cell subset within each vaccine group in the spleen (**a**, bottom) or the genital tract (**b**, bottom). Red, B cells; Green, CD8α+ T cells; Blue, CD4+ T cells; yellow, CD3- CD19- lymphocytes; Grey, remaining live CD45+ cell. Frequency (**c**, **d**) or absolute count (**e**, **f**) of each indicated cell subset in the spleen (**c**, **e**) or genital tract (**d**, **f**). Each symbol represents one animal, red horizontal bar indicates the median. Dotted vertical line separates the two vaccine groups (alum, black; Bexsero, blue) from the split of the Bexsero group into protected (dark blue) and not protected (light blue) subsets. P values are shown and were generated via Mann-Whitney analysis.

We also assessed T cell memory subsets in both tissues to determine if Bexsero vaccination impacted central or effector memory cells (Figure 3a and b). No frequency differences were noted in the spleen (Figure 3c), however there was a trend of increasing CD4+ T effector memory (Tem) cells with a corresponding trend towards decreased CD4+ naïve cells in the FGT (Figure 3d). There were no differences in the total number of cells per gram in the spleen (Figure 3e), but a significantly higher (p = 0.0263) number of CD4+ Tem in the FGT (Figure 3f). We also noted that there were significantly more (p = 0.0067) vaginal CD4+ Tcm cells per gram in protected Bexsero animals than there were in Bexsero immunized animals that were not protected (Figure 3f).

**Figure 3.**
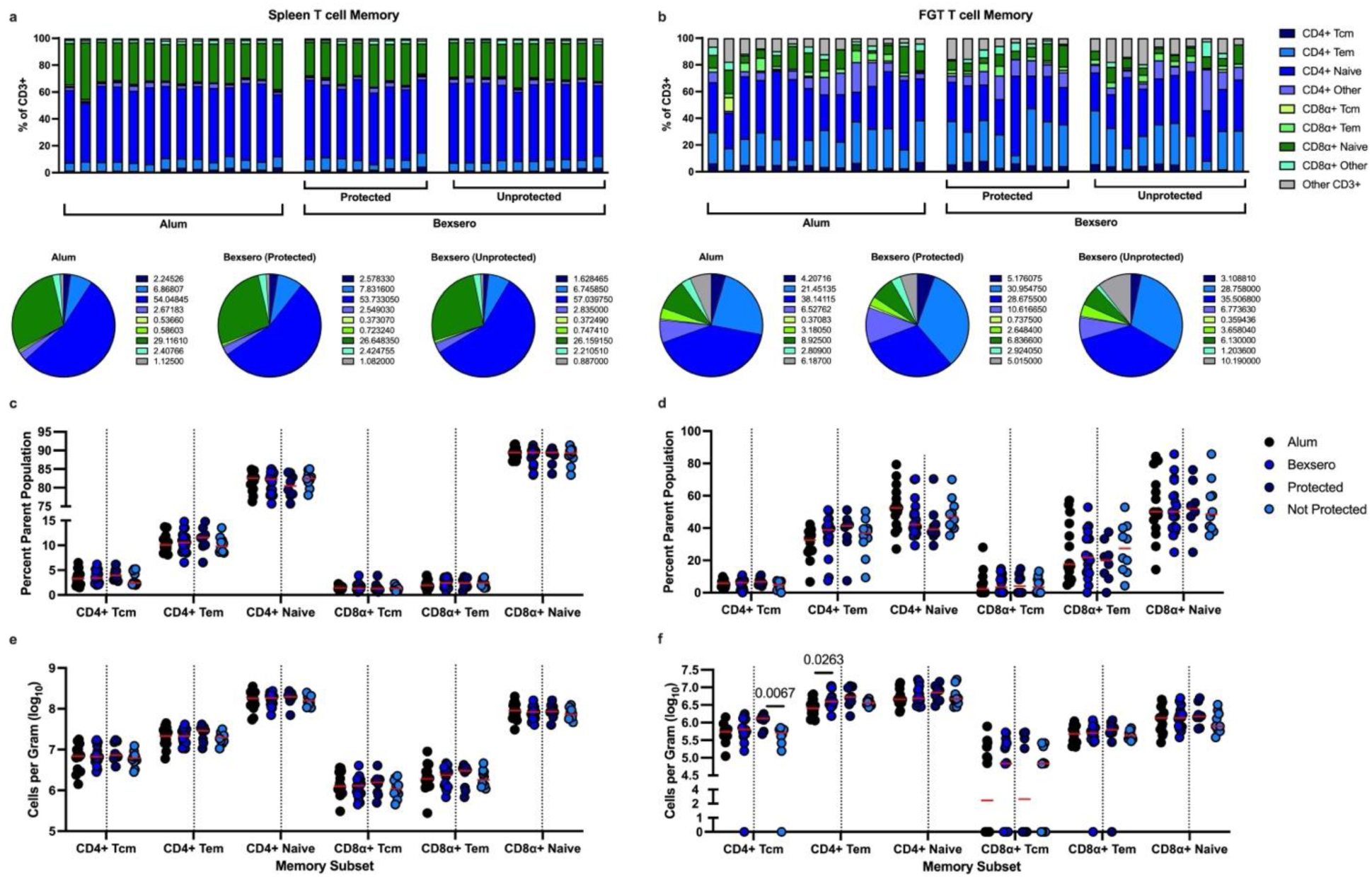
CD4+ T effector memory cell total numbers are increased in the genital tract in Bexsero vaccinated animals after infection. After three vaccinations of Bexsero or alum and six days post Ngo vaginal infection female BALB/c mice were euthanized and necropsied, and CD4+ and CD8α+ memory cell frequencies plus total counts per gram of tissue were assessed in the spleen and the female genital tract (FGT) using flow cytometry. The frequency of indicated cell subsets are shown in the spleen (**a**, Top) or the genital tract (**b**, Top), with animals divided into their vaccination group and Bexsero-protection status. Each bar represents one animal. Pie charts denote the median values of each cell subset within each vaccine group in the spleen (**a**, bottom) or the genital tract (**b**, bottom). Dark blue, CD4+ T central memory cells (Tcm; CD44+ CD62L+); light blue, CD4+ T effector memory cells (Tem; CD44+ CD62L-); blue, CD4+ naïve cells (CD44- CD62L+); light purple, all other CD4+ T cells (CD44- CD62L-); lime green, CD8α+ Tcm (CD44+ CD62L+); green, CD8α+ Tem (CD44+ CD62L-); dark green, CD8α+ naïve cells (CD44- CD62L+); light green, all other CD8α+ cells (CD44- CD62L-); grey, all other CD3+ cells. Frequency (**c**, **d**) or absolute count (**e**, **f**) of each indicated cell subset in the spleen (**c**, **e**) or genital tract (**d**, **f**). Each symbol represents one animal, red horizontal bar indicates the median. Dotted vertical line separates the two vaccine groups (alum, black; Bexsero, blue) from the split of the Bexsero group into protected (dark blue) and not protected (light blue) subsets. P values are shown and were generated by using Mann-Whitney analysis.

### CD3- CD19- lymphocytes drive a primarily T_H_1-specific immune response in Bexsero vaccinated mouse splenocytes

To uncover the vaccine-specific cytokine response generated by Bexsero immunization, splenocytes from necropsied animals were stimulated with either cell culture media alone (control) or Bexsero in the presence of co-stimulatory antibodies (anti-CD49d and anti-CD28) and a cytokine export blocking agent (brefeldin A) so individual cell responses could be measured using flow cytometry (Supplemental Figure 8). Here, adaptive responses would be considered biologically relevant if there is a difference in the media and Bexsero-stimulated samples from the Bexsero-immunized mice group and a difference in the Bexsero-stimulated samples between alum and Bexsero-immunized animals. Beyond this, the flow data sets also highlight numerous non-vaccination related effects of Bexsero stimulation, presumably due to endotoxin and/or other bacterial-derived inflammatory mediators in the vaccine (Figure 4 and Supplemental Figures 8 and 9).

**Figure 4.**
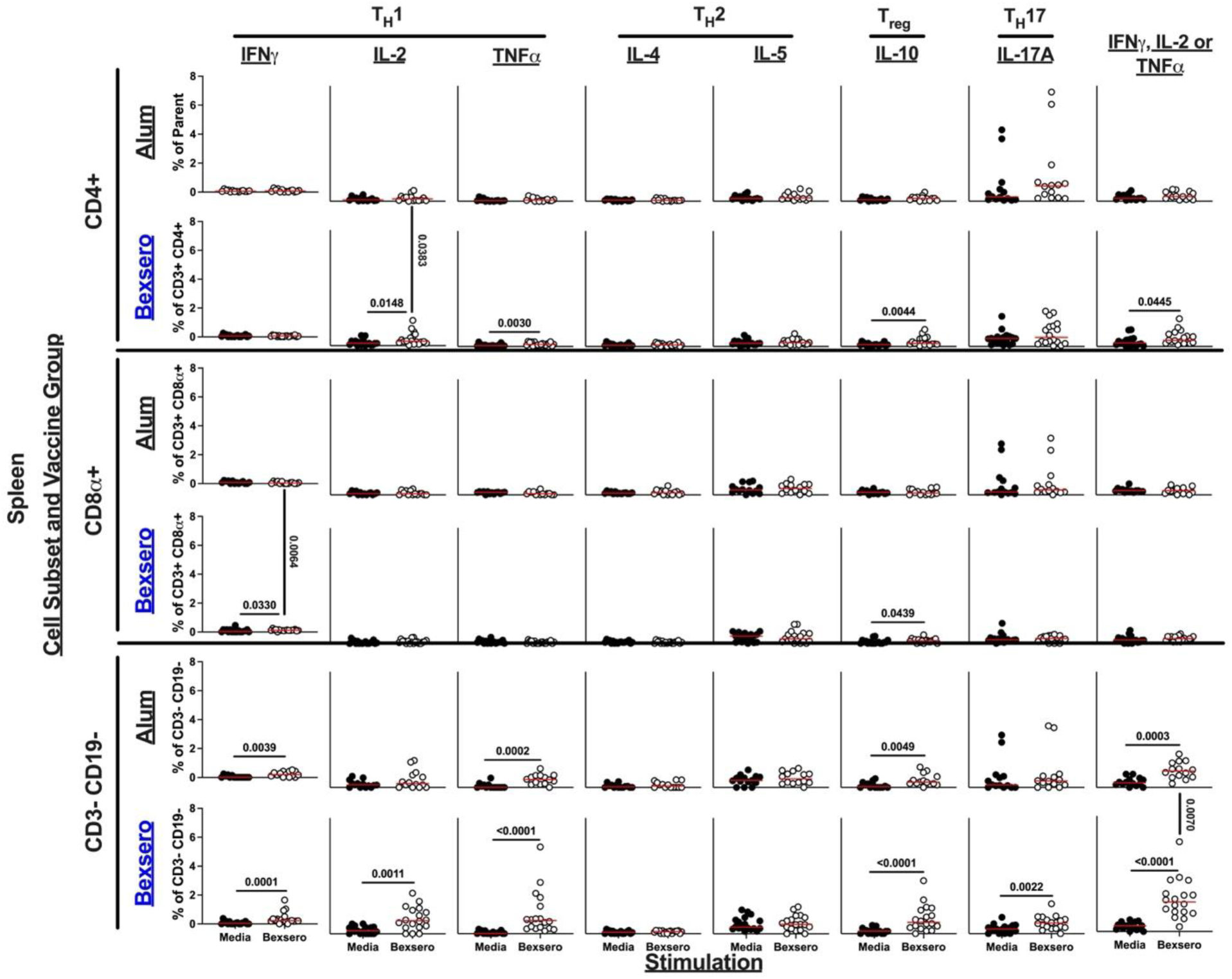
CD3- CD19- splenocytes generate a robust T_H_1 immune response in Bexsero vaccinated mice. Splenocytes from necropsied animals on day six post-infection were stimulated to assess the vaccine-specific immune response which was measured using intracellular cytokine staining and flow cytometry. Each column is the frequency measurement of a specific cytokine (or a combination of any of the T_H_1 cytokines [IFNγ, IL-2 or TNFα as performed by Boolean gating] in the far-right column) after cells were stimulated with either media alone (black) or with Bexsero (white). Rows indicate both which vaccine group is being assessed (black, alum; blue, Bexsero) and which cell subset (CD4+, CD8+, or CD3- CD19- lymphocytes). Each symbol is an animal, and the red horizontal bar represents the median of each data set. Non-parametric (Mann-Whitney) analysis was performed both within the vaccine groups (between media and Bexsero stimulation) and between groups (the alum or Bexsero stimulated cells compared between alum and Bexsero vaccinated animals). Significant p values are listed with a horizonal black bar if it was within groups or a vertical black bar if it was between vaccine groups.

Cytokines spanning numerous lymphocyte responses were analyzed including T_H_1 (IFNγ, IL-2, and TNFα), T_H_2 (IL-4 and IL-5), T regulatory (T_reg_, IL-10) and T_H_17 (IL-17A) in both conventional (CD4+ and CD8α+) and unconventional (CD3- CD19-) cell subsets (Figure 4). Our experiment demonstrated that Bexsero vaccination elicits a significant increase in the frequency of CD4+ IL- 2+ cells (p = 0.0148, media vs. Bexsero stimulation, Bexsero-vaccinated animals; p = 0.0383, alum vs. Bexsero vaccinated animals, Bexsero stimulation) and CD8α+ IFNγ+ cells (p = 0.0330, media vs. Bexsero stimulation, Bexsero-vaccinated animals; p = 0.0064, alum vs. Bexsero vaccinated animals, Bexsero stimulation). Given the importance of cells producing multiple cytokines in vaccine responses^15^, we also performed Boolean gating to capture cells producing any combination of T_H_1 cytokines (IL-2, IFNγ and TNFα). Unexpectedly, we observed a statistically significant increase in the frequency of CD3- CD19- cells producing any T_H_1 cytokine (p < 0.0001, media vs. Bexsero stimulation, Bexsero-vaccinated animals; p = 0.0070, alum vs. Bexsero vaccinated animals, Bexsero stimulation). When analyzing the responses on a number of cells per gram basis, we did not see any significantly different Bexsero-mediated immune responses, although CD3- CD19- TNFα+ cells (p <0.0001, media vs. Bexsero stimulation, Bexsero-vaccinated animals; p = 0.0504, alum vs. Bexsero vaccinated animals, Bexsero stimulation) and CD3- CD19- cells producing any T_H_1 cytokine (p = 0.0002, media vs. Bexsero stimulation, Bexsero-vaccinated animals; p = 0.0504, alum vs. Bexsero vaccinated animals, Bexsero stimulation) were approaching significance (p = 0.0504, Supplemental Figure 8). Delving into the CD3- CD19- cells producing different combinations of T_H_1 cytokines further, we noted that Bexsero-vaccinated mice had statistically higher CD3- CD19- cells producing IL-2 and TNFα only compared to control animals based both on a percent-parent population (p < 0.05) and a total cell per gram basis (p < 0.05) (Supplemental Figure 9a). Importantly, when comparing only protected from not protected Bexsero-vaccinated animals, these statistical differences held, suggesting that this cellular phenotype (robust production of TNFα with either IL-2 or IFNγ by antigen-stimulated CD3- CD19- cells) may serve as important performance indicator (Supplemental Figure 9b).

### Multiplex analysis demonstrates a robust cytokine response to Bexsero immunization

For a broader assessment of the vaccine-specific cytokine responses generated by Bexsero immunization, lymphocytes from the spleen and the FGT were stimulated with either Bexsero or cell culture media alone, and secreted cytokines were then measured by a T cell-focused 18-plex assay (Figure 5 and Supplemental Figure 10). Cytokine responses from the spleen showed a multi-faceted Bexsero-specific response, with significantly higher amounts of T_H_1 (IL-2, TNFα, IL-12p40), T_H_2 (IL-4, IL-5, IL-13), T_reg_ (IL-10), T_H_17 (IL-17A), more broadly associated pro-inflammatory cytokines (IL-1α, IL-1β, IL-6) and other important cytokines/chemokines (GM-CSF, KC, LIX, MCP-1 and MIP-2). Notably, T_H_2 (IL-4, IL-13) and IL-6 were significantly elevated in the Bexsero- vaccinated protected group compared to the not protected group. Significant vaccine-specific responses in the FGT were more limited and included T_H_1 (IL-2), T_H_2 (IL-4) and T_H_17 (IL-17A). In both data sets there were also numerous non-memory, inflammatory-related effects of Bexsero stimulation (Figure 5 and Supplemental Figure 10).

**Figure 5.**
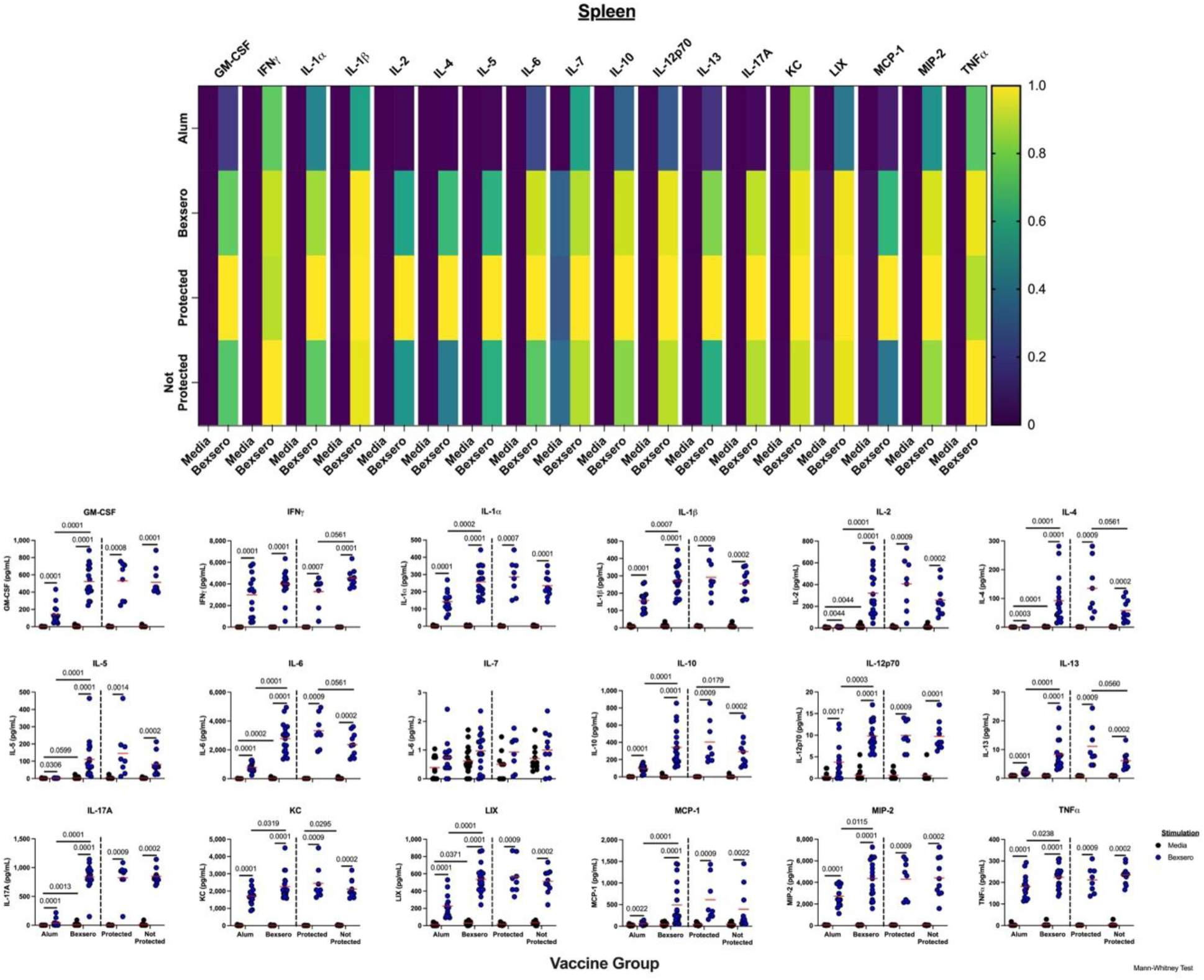
Bexsero vaccinated splenocytes secrete a multitude of cytokines from across the T cell subtype spectrum in response to Bexsero stimulation. Splenocytes from vaccinated and infected animals were incubated with either media alone or Bexsero and the secreted cytokine response from the culture media was measured using a T cell centric 18-plex. Top, a heatmap of normalized values for each cytokine listed along the top from when cells were stimulated with either media or Bexsero (columns) in the indicated group (top row, alum vaccinated animals; second row, all Bexsero vaccinated animals; third row, only protected Bexsero animals; bottom row, not protected Bexsero vaccinated animals). Bottom, Scatterplots of the raw cytokine data used to generate the heat map. Each cytokine is listed with each symbol representing the measure of an individual animal’s splenocytes after stimulation with either media (black) or Bexsero (blue). Indicated groups are listed on the X-axis. Red horizontal lines indicated the median values, and black horizontal lines connect two compared groups with statistically different medians (using Mann-Whitney non-parametric comparison) with p values listed above.

### Cytokines and chemokines in the vaginal lumen decrease as gonococci are cleared in Bexsero- vaccinated animals

To better understand the environment in the vaginal lumen prior to and throughout gonococcal infection, we assessed 45 different analytes prior to infection, as well as on days three and six post-infection. The only cytokine that was different between vaccine groups prior to infection was a statistically reduced concentration of the tissue inhibitor of metallopeptidases (TIMP-1) in the vaginal lumen of Bexsero-vaccinated animals (p = 0.0262; Supplemental Figure 12). On day three post-infection Bexsero-vaccinated protected animals had statistically lower IL-7 and erythropoietin (EPO), and some animals displayed higher IL-1β compared to animals that were not protected (p = 0.0453, 0.0354 and 0.0183, respectively; Supplemental Figure 11). Lastly, on day six we saw a reduction in twelve different analytes in Bexsero vaccinated animals, including chemokines (eotaxin, fractalkine, MCP-5, MDC, TARC), growth factors (G-CSF and M-CSF), pro-inflammatory cytokines (IL-1β, IL-6, IL-12p40, IL-16) and one cell differentiation inhibitor (LIF), whereas the only difference in the protected versus not protected Bexsero vaccinated group was less EPO in the protected group.

### Principal component analyses of complex datasets

To consider together the variability in responses among the multiple complex datasets described above, principal component analysis (PCA) was performed. For the 18-plex T cell cytokine data set from Bexsero-stimulated splenocytes 80.7% of the variability could be attributed to the first three components (Supplemental Figure 12a). The resulting loading matrix shows that the first principal component (PC1) is positively correlated with many of the measured analytes (Supplemental Figure 12b). This implies PC1 is a multifaceted measurement, driven by production of T_H_1 (IL-2, IL-12p70), T_H_2 (IL-4, IL-5, IL-13), T_H_17 (IL-17A), T_reg_ (IL-10), and other potent cytokines and chemokines (GM-CSF, IL-1 α, IL-1β, LIX, MCP-1, KC). PC2 increased with T_H_1 cytokines (TNFα and IFNγ) and decreased with T_H_2 cytokines (IL-5, IL-13). PC3 was primarily driven by positive correlation with IL-7, a growth factor that stimulates T, B and NK cells. Scores for PC1, PC2, and PC3 were compared between the vaccine groups, and it was shown the animals given Bexsero had a higher mean PC1 score compared to animals given alum (p < 0.0001, Supplemental Figure 13c), whereas PC2 and PC3 showed no difference between groups (Supplemental Figure 12d and e).

We also performed PCA on the flow cytometric frequencies (Supplemental Figure 13) and total cell counts per gram of tissue (Supplemental Figure 14) of Bexsero-stimulated splenocytes, as well as on the 45-plex cytokine data from the vaginal lavage of animals pre- and post-infection (Supplemental Figures 15). Notable highlights from these analyses included significantly higher scores in principal components from Bexsero vaccinated animals comprised of positively correlated frequencies and total cells of CD3- CD19- cells secreting TNFα alone or in combination with other T_H_1 cytokines (Supplemental Figures 13d and 14e, respectively). We also observed a significantly lower score in a total cell principal component that was highly correlated with total CD4+ T central memory and CD8α+ cells producing cytokines (Supplemental Figure 14d). Bexsero vaccinated animals had a statistically lower score in a day six post-infection vaginal lavage component that was highly correlated with a large number of chemokines and pro-inflammatory cytokines (Supplemental Figure 15, bottom) whereas protected Bexsero immunized animals had a lower average score compared to not protected animals in a pre-infection lavage component that was positively correlated with certain chemokines (fractalkine and KC) and negatively correlated with others (TARC, 6Ckine/Exodus 2, IP-10) (Supplemental Figure 15, Top). These findings both support our previous results of the importance of CD3- CD19- cells that produce TNFα and other cytokines and provide novel directions to explore regarding vaginal cytokine and chemokine responses.

## Materials and Methods

### Animals

All animal procedures were approved by the Local Animal Care Committee of the University of Toronto (permit number 20011775), which is subject to the ethical and legal requirements under the Province of Ontario, Canada, Animals for Research Act, and those of the federal Canadian Council on Animal Care (CCAC). All efforts were made to avoid and/or minimize suffering.

Mice used in this study were six- to eight-week-old BALB/c females (Charles River). Upon reception to the University of Toronto Division of Comparative Medicine (DCM) animal facility, animals were acclimatized to the environment for at least seven days before any handling was performed. One pilot experiment and two independent vaccination and challenge studies were conducted for this study. A list of experimental animals and corresponding information can be found in Supplemental Table 1.

### Vaccination

Bexsero (4CMenB, GlaxoSmithKline) vaccine was purchased at a pharmacy and stored at 4°C until use. Mice were subcutaneously (s.c.) vaccinated with either a one-half human dose (0.25 mL) or an equivalent amount of aluminum hydroxide adjuvant (0.25 mg) plus phosphate-buffered saline (PBS; Wisent Bioproducts, Canada). Vaccinations occurred every three weeks for a total of three vaccinations, with a two-week rest period before animals were staged into infection.

### Staging and preparation of mice

To determine the stage of the reproductive cycle, mouse vaginas were lavaged daily with 30 μL of PBS using a 100 μL micropipette. Wet smears were examined microscopically under a 40x objective lens, and the stage of the estrous cycle was determined based on cytology^16^. Mice transitioning from estrus or metestrus to diestrus over a 24-hour timespan were considered cycling normally and included in the study. Upon confirmation of proper cycling, mice were s.c. injected with 200 μL water-soluble β-estradiol (0.5 mg/mouse; Sigma-Aldrich, Canada) to induce estrus two days prior to infection. Mice were subsequently injected on the day of infection and day two post-infection to sequester the mice in this stage. The cycling mice also received intraperitoneal (i.p.) injections of vancomycin (2.4 mg) and streptomycin (0.6 mg) in 200 μL of PBS for four days (day −2 to day +1 relative to infection). During the same time span, mice also received trimethoprim antibiotics in their drinking water at a final concentration of 0.4 mg mL^-1^. On day 2 post-infection, the antibiotics in the water changed to additionally contain streptomycin sulphate (2.4 mg mL^-1^). This regimen continued throughout the entirety of the experiment.

### Preparation of *Neisseria gonorrhoeae* inoculum

*Neisseria gonorrhoeae* strain FA1090 was streaked out onto GC agar (Becton Dickinson, Canada) supplemented with Kellogg’s (D-glucose 4 g/L, glutamine 50 mg/L, ferric nitrate 5 mg/L, and cocarboxylase 0.2 mg/L final concentration) and VCNT (300 μg vancomycin, 750 μg colistin, 1200 U nystatin and 500 μg trimethoprim lactate; Becton Dickinson, Canada) antibiotics and was incubated overnight at 37°C plus 5% CO_2_. The overnight lawn of gonococci was harvested into 1 mL of PBS supplemented with 0.9 mM CaCl_2_ and 0.5 mM MgCl_2_ (PBS++; Wisent Bioproducts, Canada), and the optical density at 550 nm (OD_550_) was measured to calculate the desired concentration of bacteria.

### Vaginal infection and bacterial recovery

The natural estrous cycle of the mice was monitored for approximately 5 days, and once proper cycling was confirmed, hormone and antibiotic treatments were initiated as indicated above. On day 0, approximately 1 × 10^7^ colony forming units (CFU) of mouse-passaged *N. gonorrhoeae* FA1090 bacteria (see Supplemental Table 1 for exact infection doses for each animal) in 2.5 - 5 μL PBS++ were inoculated intravaginally on non-sedated mice using a P10 pipette. For gonococcal recovery, the vagina was washed daily with 15 μL of PBS++ and serial dilutions were plated on GC agar supplemented with Kellogg’s and VCNT, which suppresses the growth of commensals while allowing recovery of *N. gonorrhoeae*. Animals were considered protected if they had no bacterial recovery on at least days four through six (therefore fulfilling the cleared criteria above).

### Cell isolation for immunological assays

On day six post-infection, mice were humanely euthanized using 100% CO_2_ exposure. Both the spleen and genital tract were sterilely excised, placed into complete media (cRPMI; RPMI-1640 [Wisent Bioproducts], supplemented with 10% fetal bovine serum, 20 mM HEPES [Gibco], 1% GlutaMAX [v/v; Gibco] 10 units mL^-1^ penicillin and 10 units mL^-1^ streptomycin;) on ice and weighed. Spleens were processed into a single cell suspension by mechanically pressing tissue through a 40 μM cell strainer. Cells were treated to remove red blood cells via incubation in 1x RBC lysis buffer (eBioscience) for five minutes in the dark at room temperature. The cells were then washed with PBS and resuspended in cRPMI for counting using a haemocytometer. Female genital tracts (FGT) were finely minced using surgical scissors and incubated for 30 minutes in digestion buffer (Hanks Balanced Salt Solution [HBSS; Gibco] supplemented with 5% FBS, 10 mM HEPES [Gibco], 20 μg mL^-1^ DNAse I and 2 mg mL^-1^ collagenase D) at 37°C with shaking. The tissue was then pressed through a 40 μM cell strainer, washed with PBS and resuspended in cRPMI for counting using a haemocytometer.

### Flow cytometry

One million cells from either the spleen or the FGT were added to the wells of a round bottom 96 well plate. They were then stimulated with either a solution of Bexsero (final concentration of 0.5 μg mL^-1^ of the outer membrane vesicle [OMV; measured as amount of total protein]) and anti- CD49d (clone R1-2; final concentration 1 ug mL^-1^) plus anti-CD28 (clone 37-51; final concentration 1 ug mL^-1^) antibodies or cRPMI media with the same antibody cocktail for two hours at 37°C plus 5% CO_2_. To assess intracellular cytokines, brefeldin A (GolgiPlug, BD Biosciences) was added as per the manufacturer’s instructions, and cells were left to incubate as indicated previously for an additional 12 hours. Treated samples were then transferred to a 96-well V-bottom plate, spun at 500 × g, and washed twice with PBS. All the following steps were completed in a low to no light setting. Cells were incubated with a viability stain (BD Horizon Fixable Viability Stain 620; BD Biosciences) for 15 minutes at room temperature, washed twice with fluorescence activated cell sorting (FACS) buffer (PBS + 1% FBS), and incubated with Fc block (TruStain FcX PLUS [anti-mouse CD16/32], BioLegend) for 10 minutes at 4°C. The surface stain antibody cocktail was then added, and samples were incubated for an additional 30 minutes at 4°C. Cells were washed twice with FACS buffer and resuspended in fixation/permeabilization solution (BD Biosciences) for 20 minutes at 4°C. Samples were washed twice with Perm/Wash buffer (BD Biosciences) and the intracellular antibody cocktail was added and incubated at 4°C for 30 minutes. Samples were washed two more times with Perm/Wash buffer, resuspended in FACS buffer, and kept at 4°C in the dark until they were read on a FACSymphony™ A3 5-Laser Cell Analyser (BD Biosciences). Flow cytometry data was analyzed using FlowJo version 10.9.0 (Becton Dickinson). A list of all flow cytometry antibodies used in this study can be found in Table 1.

**Table 1.** Flow cytometry antibodies used in this study.

| **Table 1.** Flow cytometry antibodies used in this study |  |  |  |  |
| --- | --- | --- | --- | --- |
| **Marker** | **Fluorophore** | **Clone** | **Company** | **Catalogue #** |
| **Surface Panel** |  |  |  |  |
| CD3 | APC Cy7 | 17A2 | BD Pharmingen | 560590 |
|  |  |  | BioLegend | 100222 |
| CD4 | R718 | RM4-5 | BD Horizon | 566939 |
| CD19 | PerCP Cy5.5 | 1D3 | BD Pharmingen | 551001 |
| CD45 | BUV395 | 30-F11 | BD Horizon | 564279 |
| CD44 | BV786 | IM7 | BD Horizon | 563736 |
| CD62L | BUV737 | MEL-14 | BD Horizon | 612833 |
| CD8α | BUV805 | 53-6.7 | BD Horizon | 612898 |
| **Viability Stain** |  |  |  |  |
| Fixable Viability Stain 620 | NA | NA | BD Horizon | 564996 |
| **Intracellular Panel** |  |  |  |  |
| CD69 | PE Cy7 | H1.2F3 | BD Pharmingen | 552879 |
| IFNγ | APC | XMG1.2 | BD Pharmingen | 554413 |
| IL-10 | BV421 | JES5-16E3 | BD Horizon | 563276 |
| IL-17A | BV510 | TC11-18H10 | BD Horizon | 564168 |
|  |  |  | BioLegend | 506933 |
| IL-4 | BV605 | 11B11 | BD Horizon | 564007 |
| TNFα | BV650 | MP6-XT22 | BD Horizon | 563943 |
| IL-5 | PE | TRFK5 | BD Pharmingen | 554395 |
| IL-2 | FITC | JES6-5H4 | BD Pharmingen | 554427 |
| **Co-Stimulatory Antibodies** |  |  |  |  |
| anti-CD49d | NA | R1-2 | BD Pharmingen | 553153 |
| anti-CD28 | NA | 37.51 | BioLegend | 102115 |

### Cytokine multiplex assays

One million cells from either the spleen or the FGT were added to the wells of a round bottom 96 well plate and were stimulated with either Bexsero (final concentration of 0.5 μg mL^-1^ of the OMV [measured as amount of total protein]) or cRPMI media alone for 24 hours at 37°C plus 5% CO_2_. Cells were then spun down at 500 x g, and supernatants were stored at −20°C. Murine vaginal lavage samples from two weeks post-vaccination (pre-infection) as well as days three and six post-infection were spun down, and supernatants were heat treated at 56°C for 30 minutes (to kill any live *N. gonorrhoeae*) and subsequently stored at −20°C. All samples were shipped on dry ice to Eve Technologies (Calgary, Alberta, Canada) to perform multiplex assays.

### *Neisseria*-specific antibody titer analysis

Nme NZ98/254 and Ngo FA1090 were grown overnight on GC-Kellogg’s at 37°C plus 5% CO_2._ Lawns were resuspended in PBS++ and OD_600_ measured and adjusted to an OD = 0.5 by adding additional PBS++. Bacterial suspensions were heat-inactivated at 57°C for 1 hour and 20 μL/well was used to coat 384-well ELISA plates (MaxiSorp, Non-Sterile, ThermoFisher, cat. 464718) and dried in a biological safety cabinet. Once dried, plates were stored at 4°C until use. Plates were blocked with 5% BSA for 2 hours at room temperature, washed 3 times with PBS plus 0.05% Tween (PBST), incubated overnight at 4°C with mouse serum diluted in 1:20,000 1% BSA or vaginal lavages diluted 1:5 or 1:2 in 1% BSA, washed again 3 times with PBST, incubated with secondary detection antibody diluted in 1% BSA for 2 hours at room temperature. For isotypes, alkaline phosphatase conjugated goat-anti-mouse IgG1, IgG2a, IgG2b, IgG3 antibodies (Jackson ImmunoResearch, cat. 105-055-205, 105-055-206, 105-055-207, 105-055-209 respectively) at 1:5,000 dilutions were used, and plates subsequently washed and developed with BluePhos substrate (Mandel KP 5120-0059) and reads obtained at 620nm. For isotype comparison, plates were set up to allow for all 4 isotypes to be quantified for all samples against both Nme and Ngo on a single plate. Reads from 3 replicate plates were averaged. For total IgG and IgA quantification, goat anti-mouse secondary detection antibody conjugated to HRP was diluted 1:10,000 for IgG (peroxidase conjugated goat anti-mouse IgG H&L, Jackson ImmunoResearch, cat. 115-035-003) and 1:5,000 for IgA (peroxidase conjugated goat anti-mouse IgA, Southern Biotech, cat. 1040-05). Some wells of the same plates were coated with anti-mouse-IgG/IgA capture antibody instead of heat-inactivated bacteria, and standards ranging from 125 ng/mL to 1.953 ng/mL were added to these wells at the same time as the primary. ELISAs with peroxidase-conjugated secondary antibodies were developed with KPL SureBlue TMB Microwell Peroxidase Substrate (SeraCare, cat. 5120-0077), reactions were quenched with 2N H_2_SO_4_, and readings were obtained at 450/520 nm.

### Western blots

Nme NZ98/254 and Ngo FA1090 were grown overnight on GC-Kellogg’s at 37°C, 5% CO_2._ Lawns were resuspended in PBS++ and OD_600_ measured and adjusted to 3. Bacterial suspension was added to 2x sodium dodecyl sulphate (SDS) buffer (0.125 M Tris, 4% (w/v) SDS, 20% (v/v) Glycerol and 0.01% (w/v) bromophenol blue) at a ratio of 1:2, boiled for 10 minutes and then stored at −20°C until use. 2-mercaptoethanol was added to the sample at a final concentration of 5%, 25 µL sample/well was resolved on 10% SDS-PAGE. After transfer, PVDF membranes were blocked in 5% skim milk in Tris buffered saline with 0.05% Tween (TBST) for 2 hours at room temperature, and then incubated overnight at 4°C with 1:5,000 dilution of terminal mouse serum in 1% skim milk. After 3 TBST washes, peroxidase conjugated goat-anti-mouse IgG H+L (Jackson Immunoresearch, cat no. 115-035-003) was used at 1:20,000 dilution in 1% skim milk for 2 hours at room temperature, wash 3 times with TBST and developed with the Novex™ ECL Chemiluminescent Substrate Reagent Kit (Invitrogen). Images were obtained with an Invitrogen iBright CL750 Imaging System and densitometry analysis performed on ImageJ software (v1.54g).

### Serum bactericidal assay

Ngo FA1090 was grown overnight on GC-Kellogg’s at 37°C plus 5% CO_2_ and subcultured onto fresh media and grown for 4.5 hours in the same conditions. Colonies were resuspended in RPMI, OD_550_ measured and ∼250-1000 CFU were added to heat-inactivated mouse sera serially diluted in RPMI in 96 well round bottomed plates. After 5 minutes, either pooled normal human serum (Sigma, cat. H4522, Lot #SLCD1946) or baby rabbit complement (Cedarlane, CL3441-S100) at a final concentration of 15% and 2.5% respectively in a total reaction volume of 40 µL. After 60 minutes of incubation at 37°C, 5% CO_2_, 10 µL was plated. Serum bactericidal assay (SBA) titers were reported as the highest dilution at which there was >50% killing compared to complement source alone.

### Principal component analysis of cellular, cytokine and chemokine parameters

Principal Components Analysis (PCA) was performed on multi-dimensional data in JMP Pro version 17.2.0 (SAS Institute Inc., Cary, NC). For 18-plex T cell cytokine/chemokine data, PCA was run on Bexsero-stimulated splenocytes only. For flow cytometry data, PCA was run on both frequencies and counts from Bexsero-stimulated splenoctyes. For vaginal lavage 45-plex data, PCA was run separately at each time point. PCA was performed on correlations (of centered and standardized variables) using row-wise variance estimation. Loading matrices for the first three principal components are included in supplemental material (Supplemental Tables 2 - 7).

### Statistical analysis and graphing

Power calculations were based off the percent of colonized animal in a pilot study. Sample size was determined to be a minimum of 14 animals per group providing 80% power to detect a statistical difference in colonization at day six with an alpha of 0.05.

All statistical analyses and figure creation was performed using either JMP Pro version 17.2.0 or Prism version 10.2.3 (Graphpad Software LLC., La Jolla, CA).

## Discussion

The serogroup B meningococcal vaccine Bexsero has been shown to elicit modest but impactful cross-protection against Ngo infection^12^, raising exciting questions regarding both what cross-protective antigens are targeted by this multi-valent vaccine, and what immune mechanism(s) are responsible for protection against Ngo. In considering the latter question, we describe here a comprehensive immune analysis to evaluate the humoral and cellular immune responses elicited by Bexsero upon gonococcal challenge in mice. Extending upon prior work that considered protection and the humoral elicited by Bexsero and protection in mice^13^, we now provide the first comprehensive analysis of the cellular immune response elicited by this vaccine, and consider together the cellular and humoral responses in mice in the context of protected or not protected infection outcomes. We observed that immunization with Bexsero provided a significant reduction of the duration of vaginal colonization and decrease of bacterial load of Ngo in a subset of mice, reflecting the efficacy apparent in retrospective studies in humans who have received Bexsero immunization for protection against Nme. Similar efficacy against Ngo in the murine lower genital tract model has been reported by others^13^, which reinforces the robustness of this cross-protective response. A significant strength of our study design was the use of pilot data to properly power the main study. This allowed us to adopt a rigorous statistical framework for defining potential outcomes and correlates while also allowing us capacity to broadly evaluate serology, cellular immune responses, and cytokine responses.

Prior studies have established that Bexsero immunization is able to elicit cross-reactive antibodies that bind multiple gonococcal surface components, and in some cases, also facilitate complement-mediated lysis of gonococci^7,8,13^. We also found that vaccination with Bexsero-induced serum IgG antibodies that bound both meningococci and gonococci. We extended beyond the prior reports by dissecting the specific IgG isotypes that were binding to each antigen, as well as examine if there is animal to animal variability in the components being recognized. Negative correlation between number of colonized days and cumulative CFU with IgG2b specific for Nme NZ98/254 was an unexpected finding and is provocative considering the relatively low IgG2b signals detected against Ngo FA1090 itself. However, the negative correlation between number of colonized days and two anti-FA1090 proteins of approximately 85 kDa and 65 kDa was particularly interesting given that these correspond to the sizes of the outer membrane proteins BamA and NHBA, respectively. This highlights the utility of using such a more granular approach to determine antibody correlates due to the complexity of Bexsero’s antigen composition. From a mechanistic point of view, the contribution of these antibodies during clearance in our model remains unclear since we were unable to detect anti-gonococcal antibodies in the vaginal lumen of vaccinated animals (Supplemental Figure 2). This could either be due to our route of immunization being subcutaneous, which others have found to not elicit significant anti-gonococcal antibodies in the vagina^13^ or due to our measurements being performed on post-infection terminal lavages in which anti-gonococcal antibodies may have been specifically depleted by the infecting bacteria.

The cellular immune response we detected in our animal cohorts was revealing but highly complex. From an overall immune cell subset perspective, we observed a statistically significant increase in CD4+ T cell frequencies in the spleen and CD3+ T cells and CD4+ T effector memory cells in genital tract of vaccinated animals. Even through the magnitude of these responses was modest, it is important to note that our experiment was not designed to analyze changes occurring within antigen-specific T cell subsets specifically. Therefore, this nuanced change in overall T cell population is presumably the result of a more robust response within the antigen-specific T cell subpopulations. Also worth noting is that, because the measurements can only be monitored at the experimental endpoint, this study provides a snapshot into vaccination induced changes in lymphocyte and cytokine responses at a single post-infection timepoint, with factors such as current bacterial loads and duration of colonization contributing to the local immune dynamics in each individual.

The vaccine-induced cytokine response noted here was extremely robust. Multiplex ELISA data showed elevated levels of cytokines that are associated with many prominent immune subsets (T_H_1, T_H_2, T_reg_, T_H_17) when splenocytes harvested from Bexsero-vaccinated animals were re-stimulated. Indeed, our PCA analysis indicated that over 57% of variability in this dataset was due to one principal component that was positively correlated with many of these cytokines/chemokines and mean scores were statistically higher in Bexsero-vaccinated animals. Many of the cytokines we observed in our study closely resemble those seen from human PBMCs co-cultured with gonococci^17^, reinforcing the validity of using animal models for Neisseria. We also noted that some of the cytokines that we observed to be significantly higher in Bexsero-vaccinated animals have been previously been suggested to be important in the control of gonococcal infection, including IL-17^18^ and T_H_1^19,20^ responses. While we did not find any single cytokine response to correlate with protection, it is possible that a robust, multifaceted response is what is required to protect against Ngo, and future studies should take this into account.

The flow cytometric analysis of stimulated splenocytes indicated that, while CD4+ and CD8+ T cells from Bexsero-immunized animals contribute to production of T_H_1 (IFNγ, IL-2, TNFα) and T_reg_ (IL-10) associated cytokines, by far the most note-worthy response was coming from the non-T non-B lymphocytes. These cells were producing primarily T_H_1 cytokines (IFNγ, IL-2, and TNFα), and Boolean gating showed the main difference between both the alum and Bexsero group as well as protected and not protected Bexsero vaccinated animals were a subset of these cells producing TNFα in combination with either IL-2 or IFNγ. Non-T non-B lymphocytes in this study were defined by a negative gating strategy in that they expressed neither CD3 (the T cell receptor) nor CD19 (a B cell lineage marker), which begs the question of what exactly they are? The most likely answer is NK cells or a member of the innate lymphocyte cell (ILC) lineage 1 family given their anatomical location^21,22^ and the strong T_H_1 response observed^23,24^. The role of both of these cells subsets is gonococcal infection is largely unknown^25^, however other studies have shown that vaccines can stimulate these cell subsets and they can be critical for protection in other infection models^26,27^. Since T cells that produce more than one cytokine are important in vaccine studies^15,28^, future analyses should aim to identify this cell population and consider its ability to promote gonococcal clearance from the female genital tract.

Examining the local immune responses within the genital tract offered further insights into the complex and dynamic interplay of cytokines and chemokines over the course of infection. The unbiased analysis of 45 different cytokines on day six post-infection of vaginal lavages showed Bexsero-vaccinated animals having lower concentrations of numerous pro-inflammatory cytokines/chemokines, including IL-6, IL-12p40, IL-16, Eotaxin, MCP-5 and Fractalkine. This finding was also supported in PCA analysis of this dataset whereby Bexsero-vaccinated animals had statistically lower mean scores of a component that explained nearly 40% of the variance in this dataset and was comprised of many of the factors listed above. This appears to reflect the waning response as Bexsero-vaccinated animals clear their infection, whereas control animals would still be actively inflamed from the presence of the gonococci. Assessing vaginal lavages prior to infection were more interesting. Although only TIMP-1 was statistically lower in Bexsero-vaccinated animals compared to our controls from a concentration perspective, PCA analysis indicated protected Bexsero-vaccinated animals had statistically lower mean scores in one component which described over 10% of the variance and was negatively correlated with a few key T cell/NK cell chemoattractants (TARC, 6Ckine/Exodus 2, IP-10)^29–31^. This, along with the higher T effector memory cell counts observed in the genital tract of Bexsero-immunized animals support the premise that T cells are recruited and/or proliferate in the FGT during infection. *Ex vivo* stimulation of lymphocytes derived from the genital tract further support that vaccinated animals are capable of producing significantly greater levels of cytokines upon antigen encounter, including cytokines that promote T cell proliferation and survival (IL-2), B cell differentiation (IL-4) and mediate proinflammatory responses (IL-17), of which IL-17 has been previously linked to gonococcal clearance^18^ (reviewed by Raphael *et. al.*^32^).

The female mouse lower genital tract infection model used in this study is now widely accepted for preclinical evaluation of vaccines and antibiotics against gonorrhea, however we remain mindful of its limitations. First and foremost, the gonococci are highly adapted to life in humans, so virulence factors involved in processes such as cellular adherence and nutrient acquisition are not considered in wild type mice. While humanized transgenic mice are available to allow these factors to be considered^33,34^, the complexities associated with breeding these animals is restrictive in studies where a large cohort of female mice of the same age are required. Considering the female genital tract itself, hormonal fluctuations can influence the immune response and susceptibility to infection^35^. Given that mice in diestrus tend to resist gonococcal infection^36^, it remains unknown how the response to infection would be affected during progesterone-dominant stages of reproductive cycling. It is also worth mentioning that for our *ex vivo* antigen-stimulation experiments, we used Bexsero as the stimulant in efforts to achieve the maximum response. While gonococci and meningococci share substantial homology^37^, some key components in Bexsero such as PorA, which is a major component of the OMV, fHbp, NadA, and both fusion proteins (GNA1030 and GNA2091) are not present/surface exposed in *N. gonorrhoeae*. Thus, OMVs from diverse gonococcal isolates would provide a more accurate measure of the strain-specific diversity of antigen driven cellular responses for future correlation studies.

In summary, we have established that a robust cellular response develops following Bexsero immunization, with T effector cells and unconventional lymphocytes (non-T non-B) appearing to play a central potential role in driving gonococcal clearance. Our findings highlight that focusing solely on antibody titers and serum bactericidal assays may not reveal the mechanistic determinants of protection and that a comprehensive analysis that integrates the humoral and cellular immune responses will likely be required to define the correlates of protection. As multiple clinical trials are ongoing with Bexsero, and others are being implemented to test other vaccine formulations, we hope that insights gained from this study will benefit future vaccine assessments.

## Author contributions

JJZ developed the experimental outline, performed animal immunizations, infections, bacterial enumeration and necropsies, performed all flow cytometric analysis, statistical analysis and wrote the the manuscript. JEF developed the experimental outline, performed animal infections, bacterial enumeration and necropsies, performed antibody titer assays and wrote the manuscript. PM performed principal component analysis and wrote the manuscript. EAI developed the experimental outline, performed animal immunizations, infections, bacterial enumeration, antibody isotype assays, western blots and serum bactericidal assays, statistical analysis and wrote the manuscript. ISL performed bacterial enumeration, antibody titer analysis and edited the manuscript. CP assisted with antibody titer analysis and edited the manuscript. LC performed bacterial enumeration and edited the manuscript. SGO provided conceptuilization support, supervision and edited the manuscript.

## Acknowledgements

The authors would like to thank the animal support staff at the Division of Comparative Medicine at the University of Toronto for technical and welfare support for the animal studies presented here. We would also like to thank Dr. Nelly Leung for general laboratory support, Dr. Elissa Currie for immunization of some animals in the pilot study and all members of the Gray-Owen laboratory for generating thoughtful discussion on the data. This research was supported by Canadian Institute for Health Research grant PJT-153177 and National Institutes of Health grant 1U19AI144182-01. S.D.G. is supported by the Canada Research Chair Program.

