## supplemental figures for "Meningococcal vaccine Bexsero elicits a robust cellular immune response that targets but is not consistently protective against *Neisseria gonorrhoeae* during murine vaginal infection"

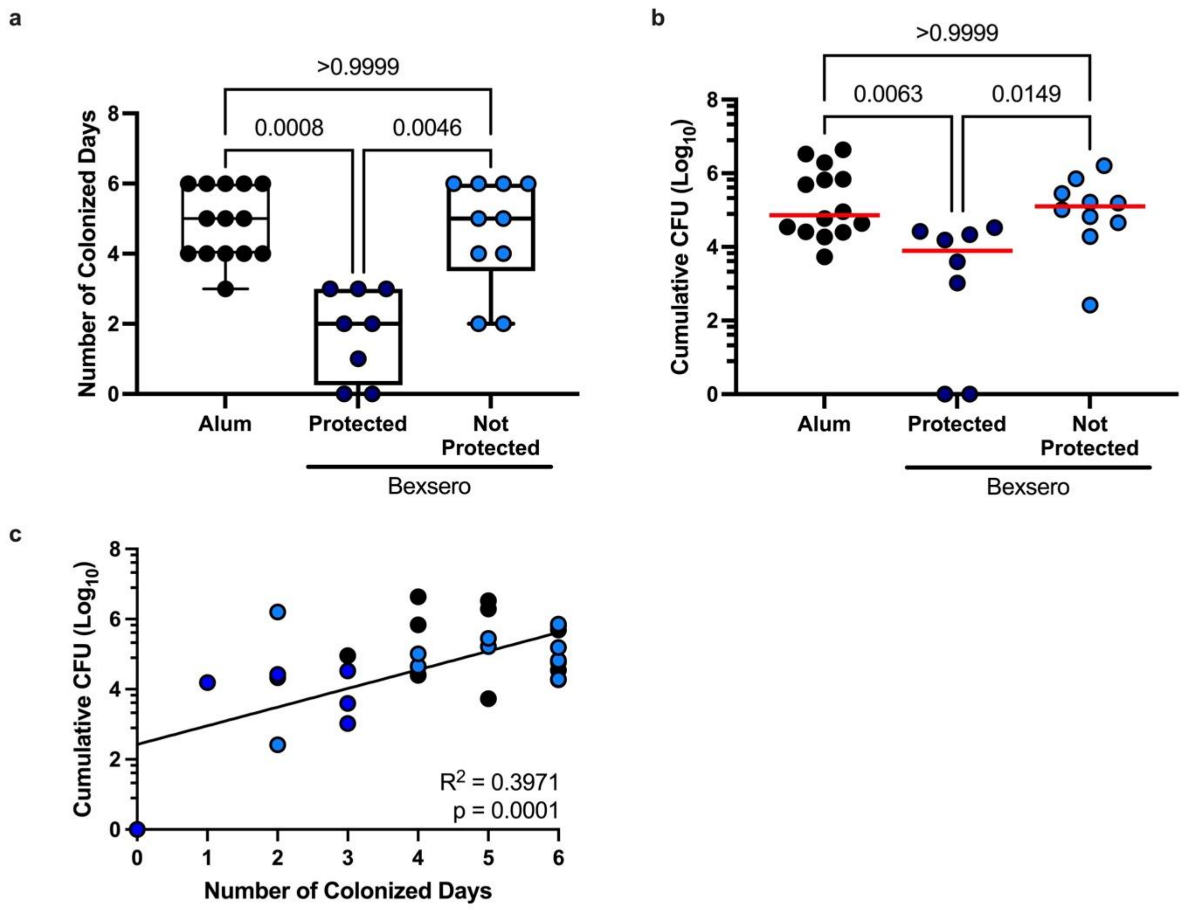


**<u>Supplemental Figure 1</u>. Comparison of colonization rates and cumulative CFU between alum controls with protected and not protected Bexsero-vaccinated animals**. Bexsero vaccinated animals were separated into protected (dark blue) and not protected (light blue) based off whether any amount of Ngo was recovered from vaginal lavages after day three of infection. Evaluation of number of colonized days (**a**) and cumulative CFU (**b**). Non-parametric Kruskal-Wallis test was performed to compare groups in both data sets and p values are listed with red horizontal times indicating the media of each group. (**c**) Simple linear regression of number of colonized days against cumulative CFU with the R squared value listed and the p value from the F test indicating the slope of the line of best fit is significantly non-zero. Alum control animals are in black, and each symbol represents one animal.

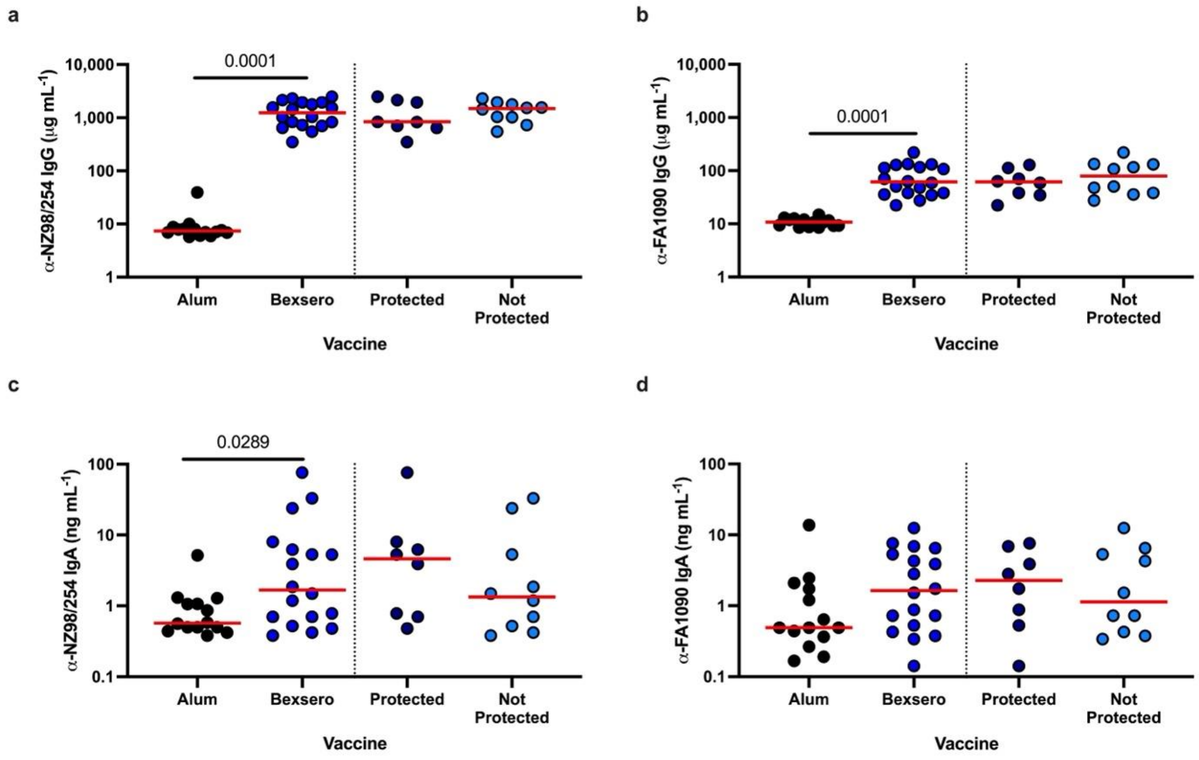


**<u>Supplemental Figure 2.</u> Bexsero immunization generates antibodies that bind *Neisseria* species.** After three vaccinations, IgG antibodies from serum (**a, b**) or IgA from vaginal lavages (**c, d**) against heat killed Nme NZ98/254 (the strain used to generate the outer membrane vesicle portion of Bexsero; **a, c**) or Ngo FA1090 (challenge strain from this study; **b, d**) were quantified. Mann-Whitney non-parametric tests were performed between alum (black) and Bexsero (blue) or protected (dark blue) and not protected (light blue) groups and significant p values are displayed. Horizontal red bars indicate the median value of each group, and each symbol indicates one animal from the study.

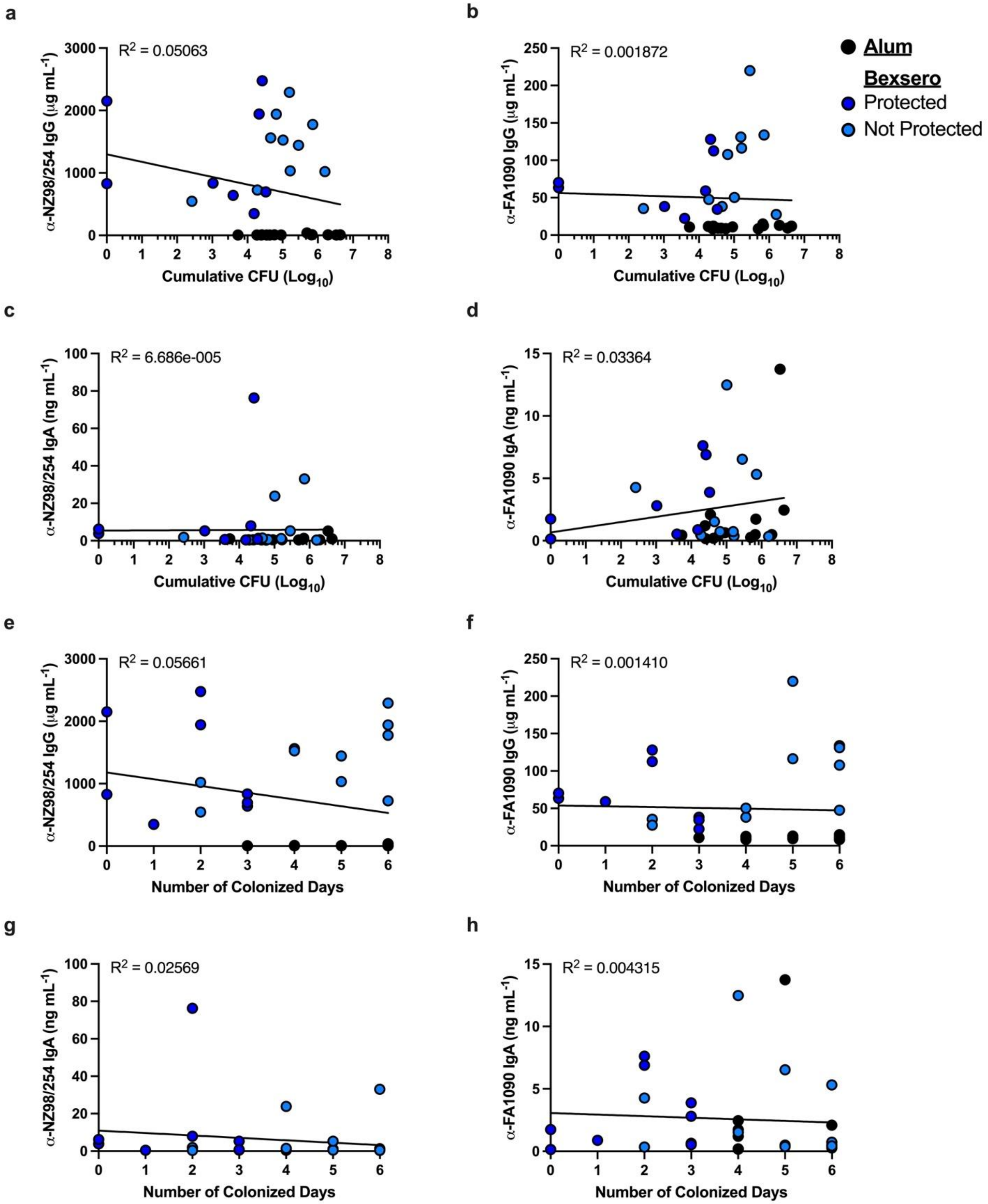

a
R2 = 0.05063
α-NZ98/254 IgG (μg mL-1)
Cumulative CFU (Log10)
b
R2 = 0.001872
α-FA1090 IgG (μg mL-1)
Cumulative CFU (Log10)
Alum
Bexsero
Protected
Not Protected
c
R2 = 6.686e-005
α-NZ98/254 IgA (ng mL-1)
Cumulative CFU (Log10)
d
R2 = 0.03364
α-FA1090 IgA (ng mL-1)
Cumulative CFU (Log10)
e
R2 = 0.05661
α-NZ98/254 IgG (μg mL-1)
Number of Colonized Days
f
R2 = 0.001410
α-FA1090 IgG (μg mL-1)
Number of Colonized Days
g
R2 = 0.02569
α-NZ98/254 IgA (ng mL-1)
Number of Colonized Days
h
R2 = 0.004315
α-FA1090 IgA (ng mL-1)
Number of Colonized Days

**Supplemental Figure 3. Anti-Neisserial antibodies do not correlate with protection.** Simple linear regression analyses were performed between serum IgG antibodies (**a, b, e, f**) or vaginal IgA antibodies (**c, d, g, h**) against heat killed Nme NZ98/254 (the strain used to generate the outer membrane vesicle portion of Bexsero; **a, c, e, g**) or *Ngo* FA1090 (challenge strain from this study; **b, d, f, h**) and cumulative bacterial CFU (**a – d**) or the number of CFU positive days (**e – h**). Each symbol represents one animal, and each group is indicated (alum, black; Bexsero protected, dark blue; Bexsero not protected, light blue). The line of best fit is present on each graph and the R squared value is listed. Statistical F tests were performed on each graph; however, no slope was significantly non-zero.

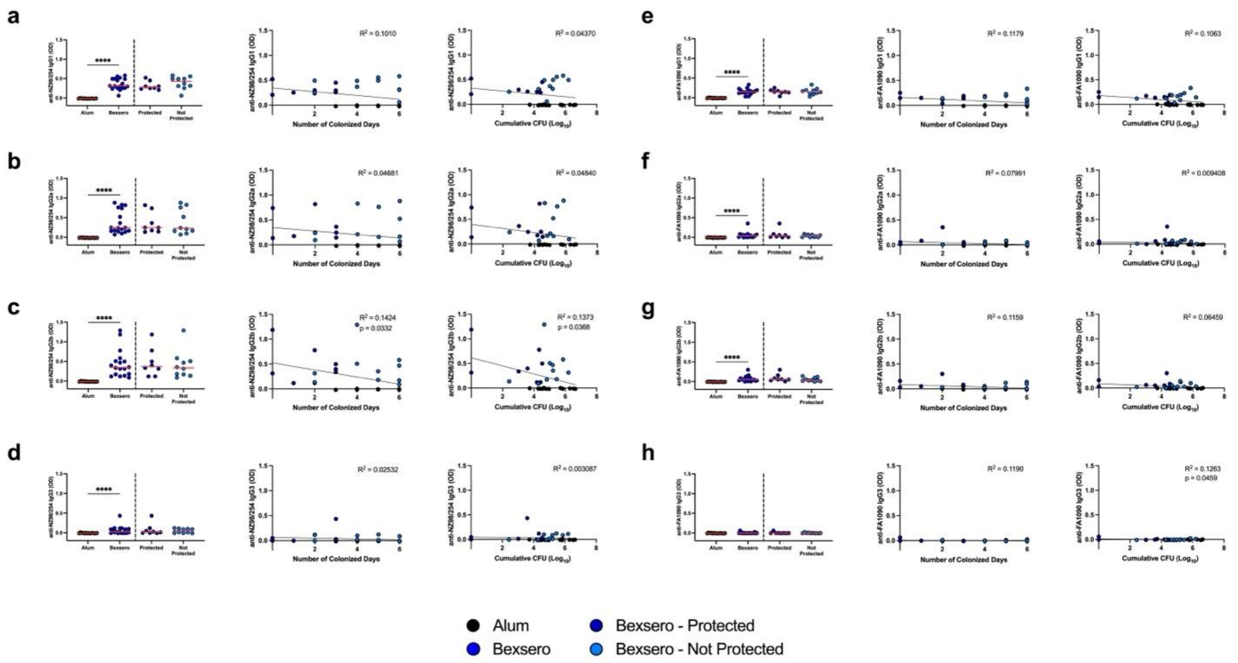


**<u>Supplemental Figure 4.</u> Distinct IgG subclasses react against *Neisseria* species following Bexsero-immunization.** IgG1, IgG2a, IgG2b, IgG3 subclasses in terminal serum of Bexsero- and alum-immunized mice against heat killed Nme NZ98/254 (**a - d**) or Ngo FA1090 (**e - h**) were measured. Mann-Whitney non-parametric tests were performed between alum (black) and Bexsero (blue) or protected (dark blue) and not protected (light blue) groups and significant p values are displayed (left panel). Horizontal red bars indicate the median value of each group, and each symbol indicates one animal from the study. Simple linear regression analyses were performed between IgG1 (**a, e**), IgG2a (**b, f**) IgG2b (**c, g**) and IgG3 (**d, h**) and the number of CFU positive days (middle panel) or cumulative bacterial CFU (right panel) is shown. Each symbol represents one animal. The line of best fit is present on each graph and the R squared value is listed. Statistical F tests were performed on each graph; and p-values for significantly non-zero slopes depicted.

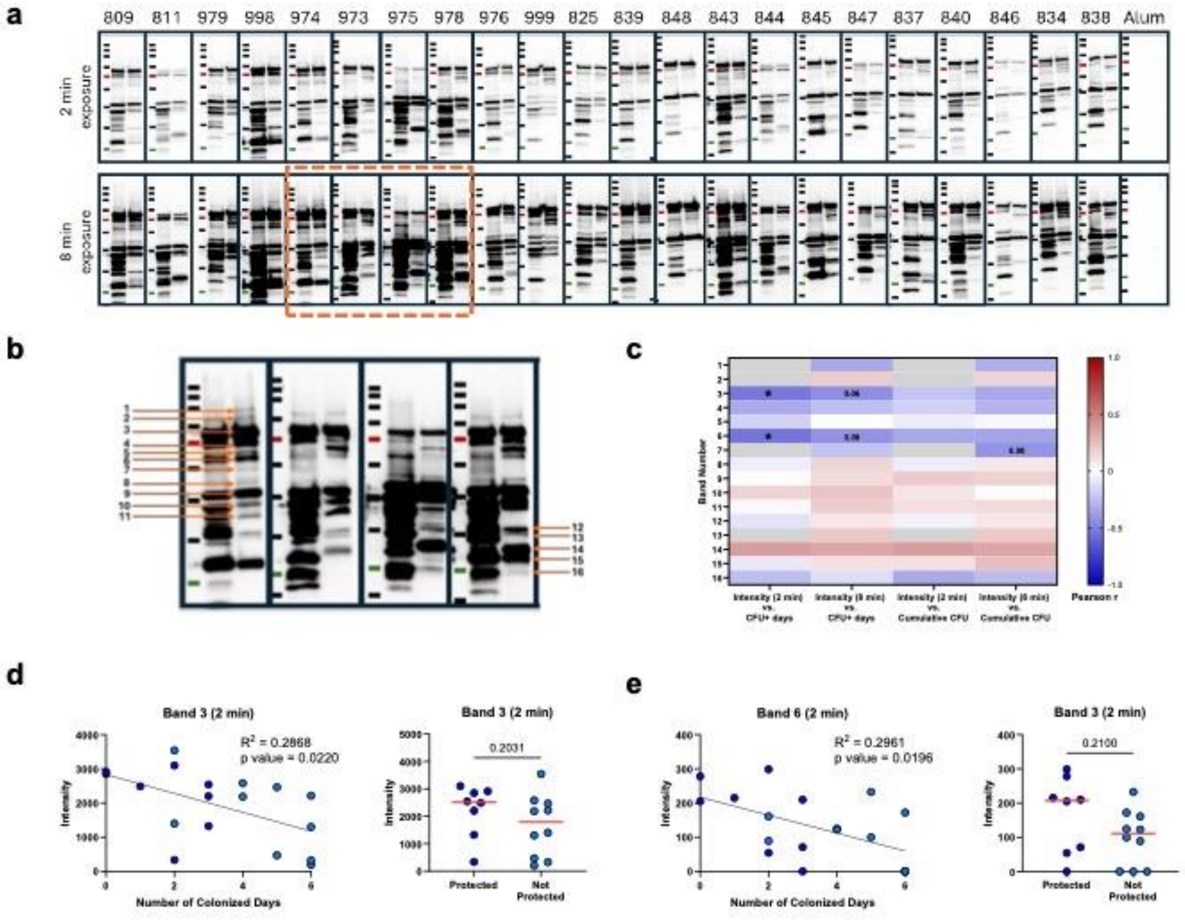


**<u>Supplemental Figure 5.</u> Variability in band recognition pattern among Bexsero-immunized animals and potential correlations.** Western blots using terminal serum from individual Bexsero-immunized mice with the identifier indicated above, and pooled sera from alum-immunized mice, against whole bacterial lysates. For each blot, lane 1 – molecular weight marker (245kDa, 180kDa, 135kDa, 100kDa, 75kDa-red, 63kDa, 48kDa, 35kDa, 25kDa-green, 20kDa, top to bottom); lane 2 - Nme NZ98/254; lane 3 – Ngo FA1090 (**a, b** – zoomed to depict individual FA1090 bands quantified by densitometry). Pearson correlation coefficient was calculated to identify potential linear relationships between intensity for each band at 2 min and 8 min exposures and number of colonized days and cumulative CFU. Pearson r values depicted as a heatmap (c); colour scale represents Pearson r values (red – positive relationship; blue – negative relationship). Interactions that could not be computed are grey. p values<0.05 indicated with *; p values <0.1 also noted. Linear regression analyses of band intensity versus number of days colonized depicted for significant interactions only, along with comparison of intensities between protected and not protected using non-parametric Mann-Whitney test (**d** – Band 3, **e** – Band 6). For scatter plots, the line regression line, $R^2$ (goodness of fit) and p values depicted.

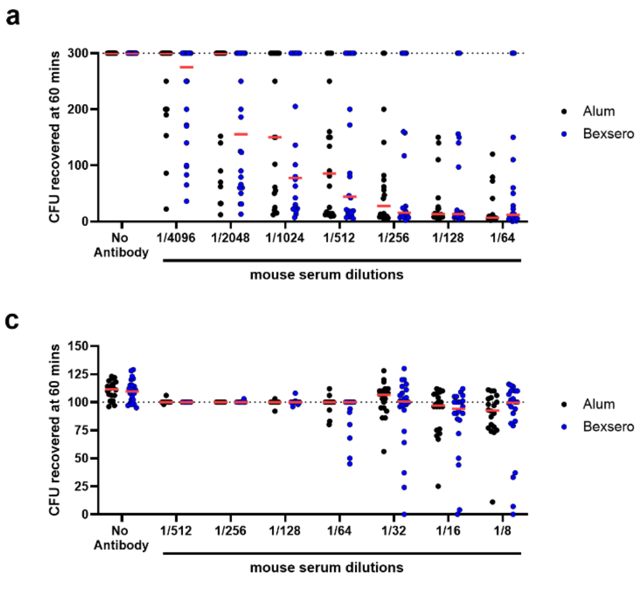


**Supplemental Figure 6. Bexsero-immunization did not augment serum bactericidal activity (SBA) titres against FA1090.** Log-phase Ngo FA1090 was incubated with increasing 2-fold dilutions of heat inactivated Bexsero or alum-immunized terminal serum and 2.5% baby rabbit complement (**a, b**) or 15% pooled human serum (**c, d**). CFU recovered after 60 minutes of incubation (**a, c**) and SBA titre, which is the highest dilution at which 50% killing was observed relative to no antibody control, (**b, d**) are graphed. Each symbol depicts serum from an individual animal; alum (black) and Bexsero (blue); Bexsero protected (dark blue) and not protected (light blue); line at median. Dotted line (**a ,c**) represents the limit for obtaining individual CFU counts; dotted lines (**b, d**) represent the highest and lowest mouse serum dilutions tested. Two-way ANOVA with Šídák's multiple comparisons test to detect differences in CFU recovered in alum versus Bexsero groups for each serum dilution (for **a**) and non-parametric Mann-Whitney test comparing alum versus Bexsero or Bexsero Protected versus Not Protected groups (for **b**) did not yield significant p values.

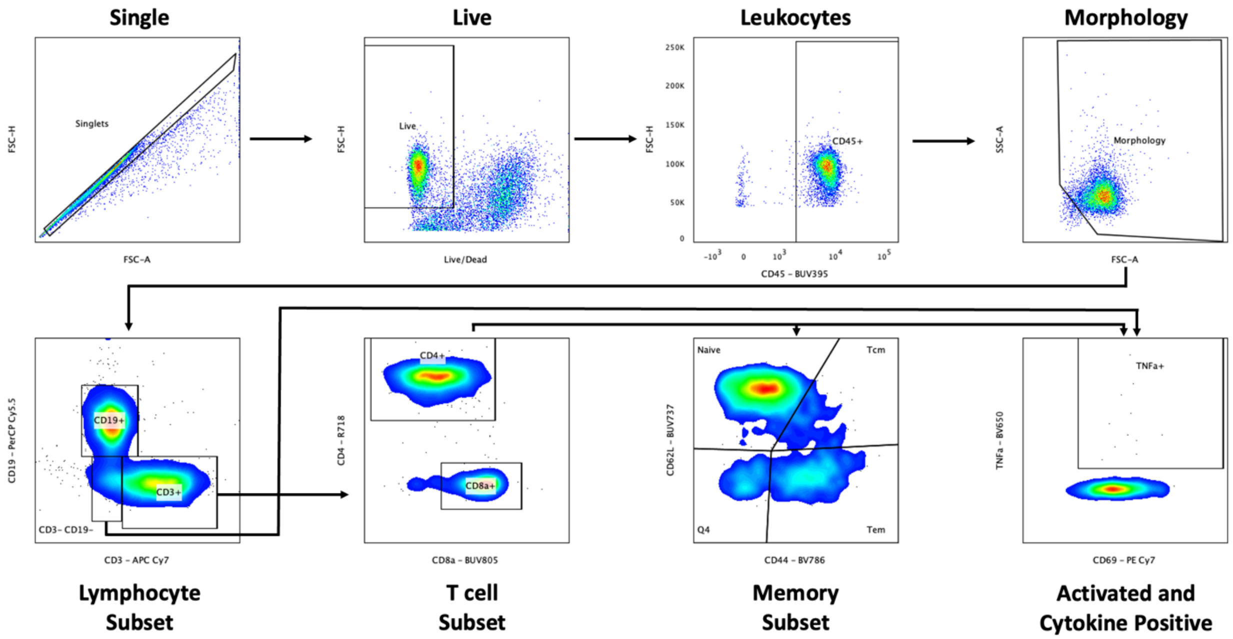


**<u>Supplemental Figure 7.</u> Flow cytometry gating strategy.** Single cell suspensions of lymphocytes from the spleen or the genital tract from vaccinated and infected animals were cultured with either media alone or Bexsero in the presence of a Golgi apparatus-blocking chemical to perform surface and intracellular flow cytometry. Cells were first gated upon as single, live (viability dye negative), leukocytes (CD45+). They were then gated on by forward and side scatter for the correct lymphocyte morphology. T cells (CD3+), B cells (CD19+) and non-T non-B lymphocytes (CD3- CD19-) were all sub-gated upon for further analysis. T cells were separated into CD4+ and CD8α+ followed by either memory markers (naïve, CD44- CD62L+; central memory [Tcm], CD44+ CD62L+; effector memory [Tem], CD44+ CD62L-) or for cytokine positivity (CD69+ plus any one of seven cytokine markers). Non-T non-B lymphocytes were gated on for cytokine positivity. Only surface gating was performed on genital tract lymphocytes.

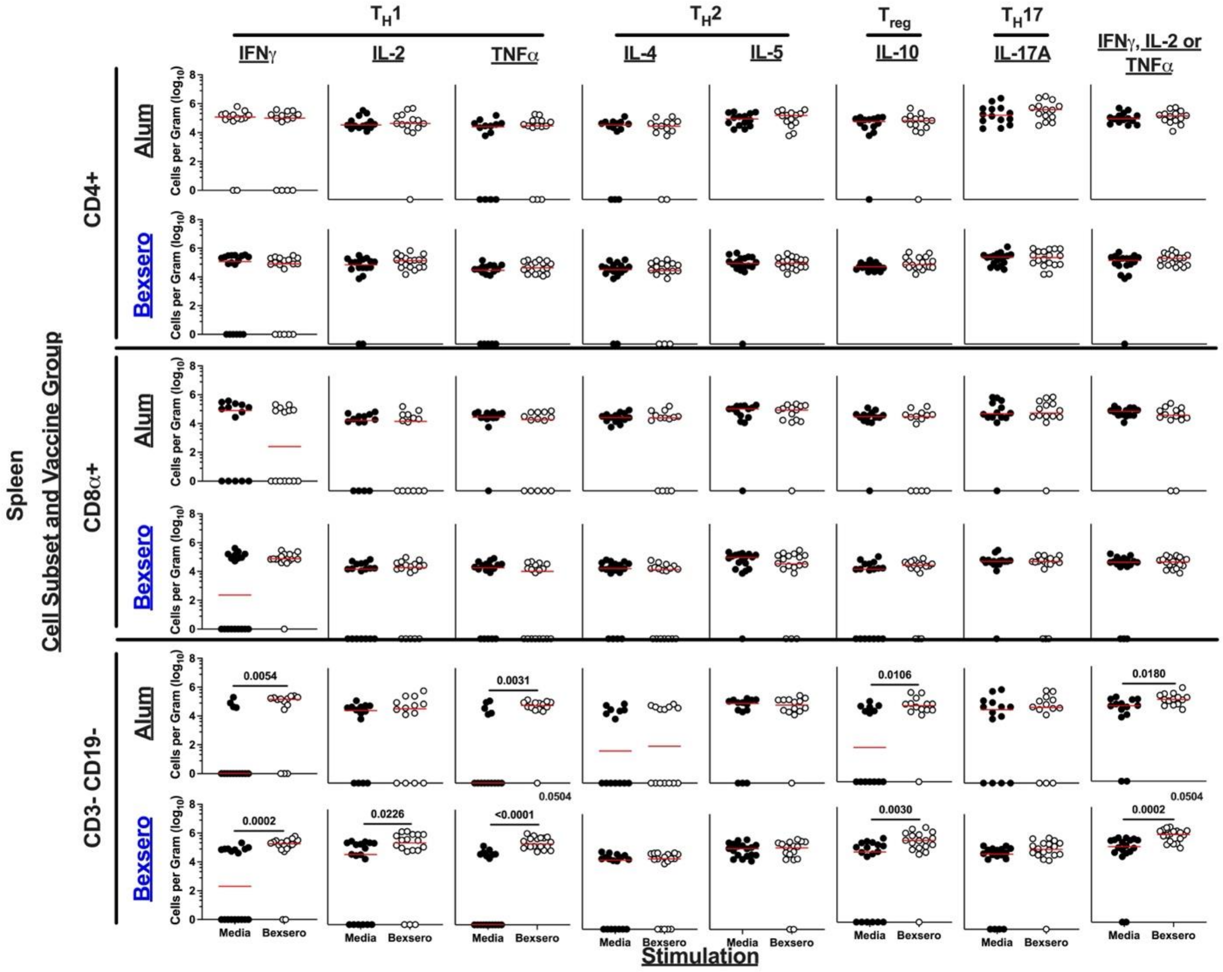


**<u>Supplemental Figure 8.</u> An increase in the total number of CD3- CD19- $T_H1$+ cells per gram is observed in Bexsero vaccinated mouse spleens.** Splenocytes from necropsied animals on day six post-infection were stimulated to assess the vaccine-specific immune response which was measured using intracellular cytokine staining and flow cytometry. Each column is the measurement of the total number of cells per gram secreting a specific cytokine (or combination of any of the $T_H1$ cytokines [IFNγ, IL-2 or TNFα] as determined by Boolean gating) after cells were stimulated with either media alone (black) or with Bexsero (white). Rows indicate both the vaccine group being assessed (black, alum; blue, Bexsero) as well as the cell subset (CD4+, CD8+ or CD3- CD19- lymphocytes). Each symbol is one animal, and the red horizontal bar represents the median of each data set. Non-parametric (Mann-Whitney) analysis was performed both within the vaccine group (between media and Bexsero stimulation) and between groups (the alum- or Bexsero-stimulated cells compared between alum and Bexsero vaccinated animals). Significant p values are listed with a horizonal bar for within group comparison or in the of the graph for between vaccine groups.

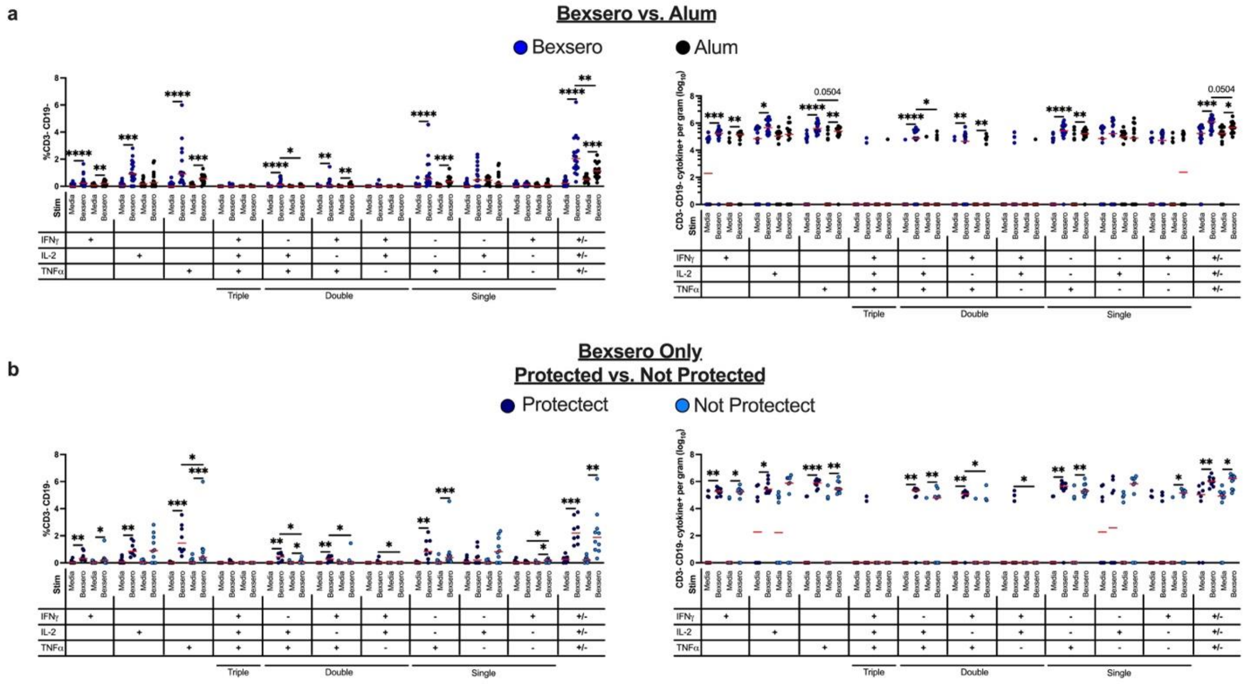


**<u>Supplemental Figure 9.</u> Boolean gating on CD3- CD19- splenocytes producing TH1 cytokines.** Splenocytes from control and Bexsero-vaccinated animals were stimulated with media alone or Bexsero and analyzed by Boolean gating flow cytometry to assess non-T non-B (CD3- CD19-) lymphocytes that were expressing any combination of $T_H1$ cytokines (IFNγ, IL-2 and/or TNFα). (**a**) Comparison of alum (black) versus all Bexsero-vaccinated (blue) animals by percent (left) or cells per gram of tissue (right). (**b**) Comparison of protected (dark blue) versus not protected (light blue) Bexsero-vaccinated animals by percent (left) or cells per gram of tissue (right). Non-parametric (Mann-Whitney) analysis was performed both within the groups (between media and Bexsero stimulation) and between groups (the alum- or Bexsero-stimulated cells compared between alum and Bexsero vaccinated animals or protected and not protected Bexsero-vaccinated animals). *, $p < 0.05$; **, $p < 0.01$; ***, $p < 0.001$; ****, $p < 0.0001$. Horizonal red bars indicate the median of each data set and horizontal black bars with the asterisk above indicate the groups being statistically compared as indicated above and are significant.

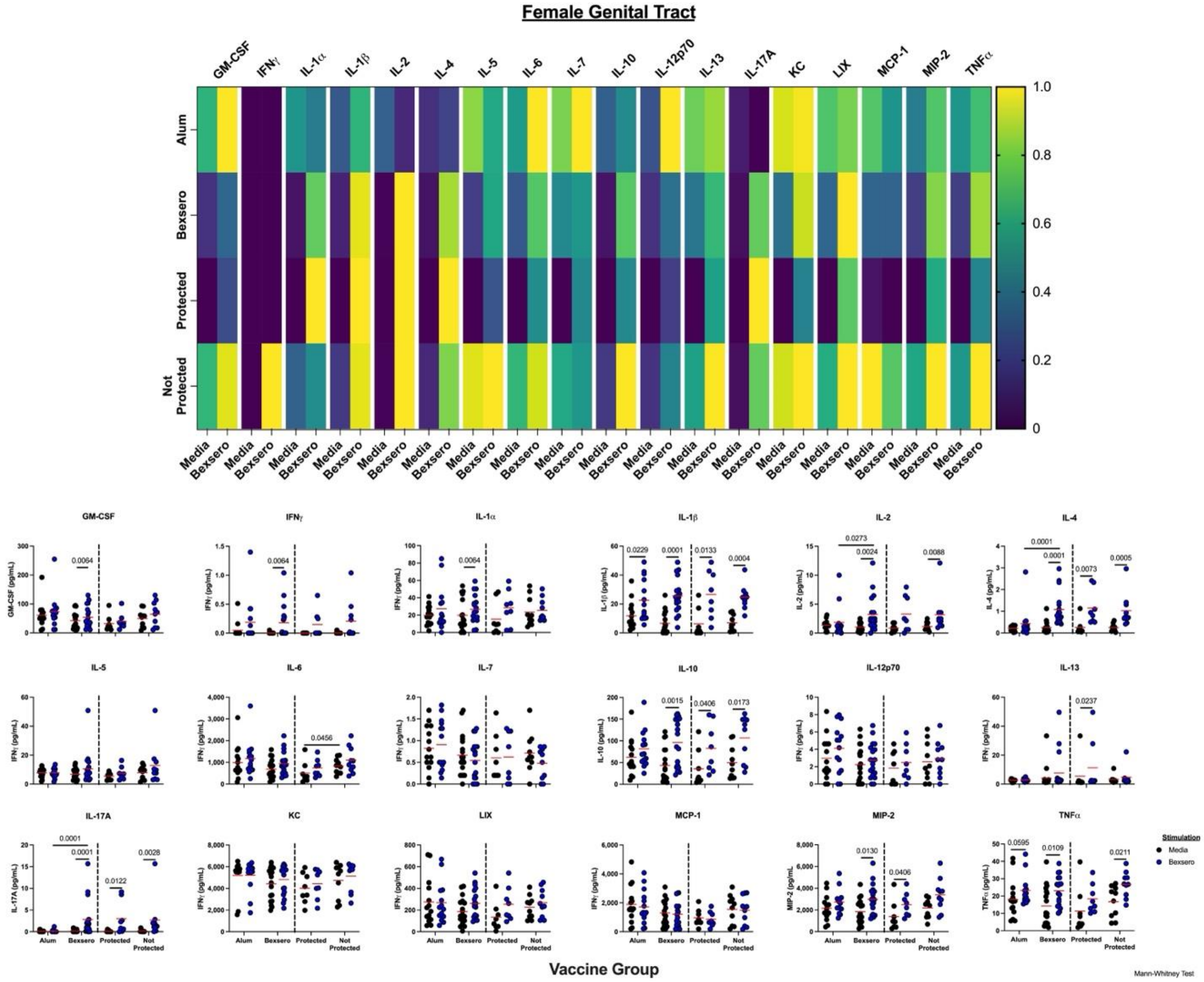


**Supplemental Figure 10.** **Lymphocytes from the genital tract of Bexsero vaccinated animals secrete more IL-2, IL-4 and IL-17A upon stimulation than their alum vaccinated counterparts.** Lymphocytes from the genital tract of vaccinated animals were incubated with either media alone or Bexsero and the secreted cytokine response from the culture media were measured using a T cell centric 18-plex. Top, a heatmap of normalized values for each cytokine listed from when cells were stimulated with the indicated agent (columns) from the specific group (top row, alum vaccinated animals; second row, all Bexsero vaccinated animals; third row, protected Bexsero animals; bottom row, not protected Bexsero vaccinated animals). Bottom, Scatterplots of the raw cytokine data used to generate the heat map. Each symbol represents the measure of an individual animal's cells after stimulation with either media (black) or Bexsero (blue). Groups are listed on the X-axis. Red horizontal lines indicated the median values, and black horizontal lines connect two groups with statistically different medians (using Mann-Whitney non-parametric comparison) with p values listed above.

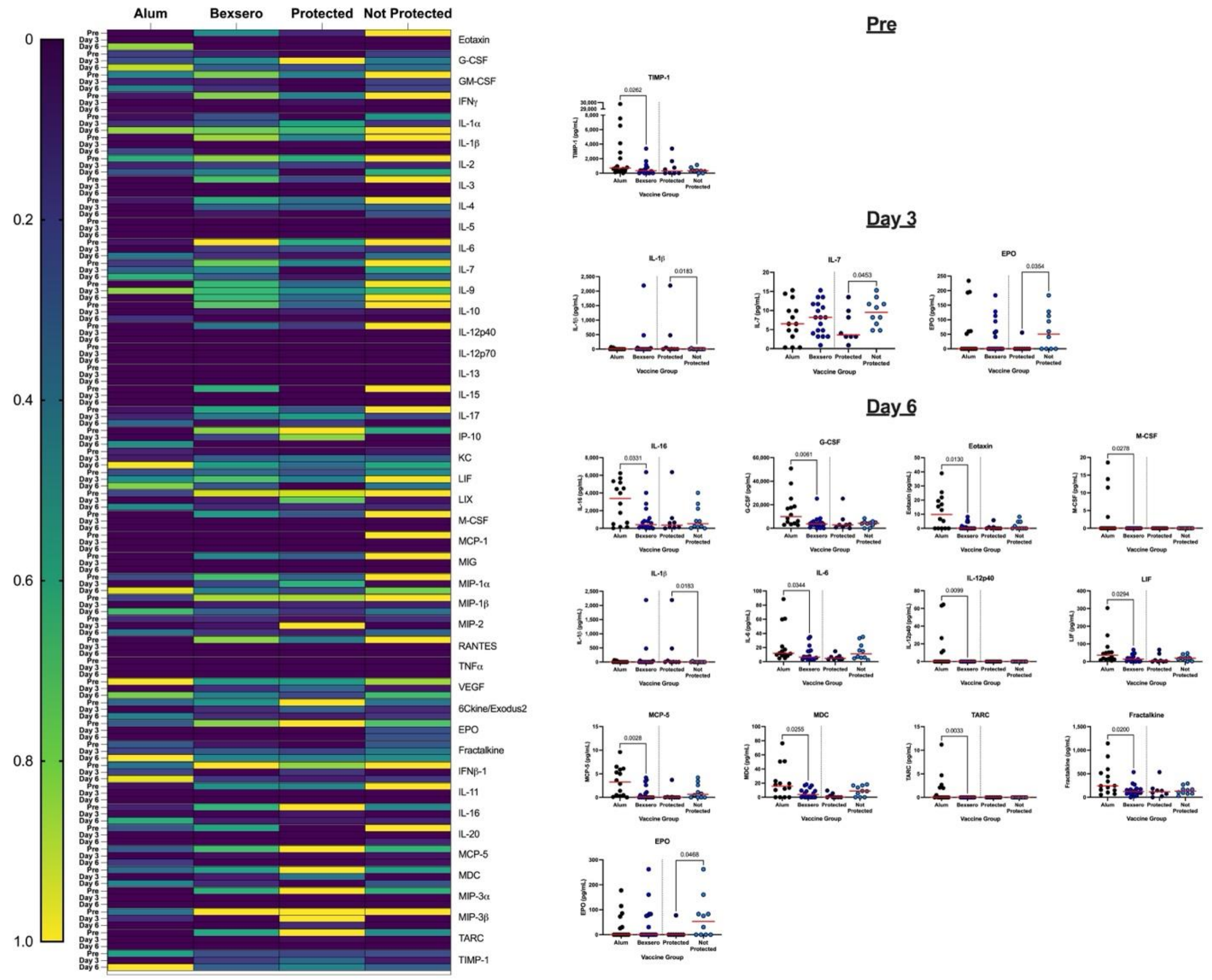


<u>Supplemental Figure 11.</u> **Multiplex analysis of vaginal lavages from prior to and during gonococcal infection.** Lavages from the vaginal lumen of all animals approximately two weeks post-vaccination (prior to infection) and days three- and six post-infection were assayed for 45 different cytokines using a multiplex assay. Left, a heat map of the normalized median values for each cytokine (indicated at the left) on each day (indicated on the left) in each specific group (indicated on the top). Right, scatter plots of all cytokines that were statistically different between the Bexsero and alum vaccinated groups or protected or not protected Bexsero-vaccinated groups as determined by Mann-Whitney non-parametric comparison. Each symbol is one animal (black, alum; blue, Bexsero; protected Bexsero, dark blue; not protected Bexsero, light blue) and red horizontal bars indicate the median value.

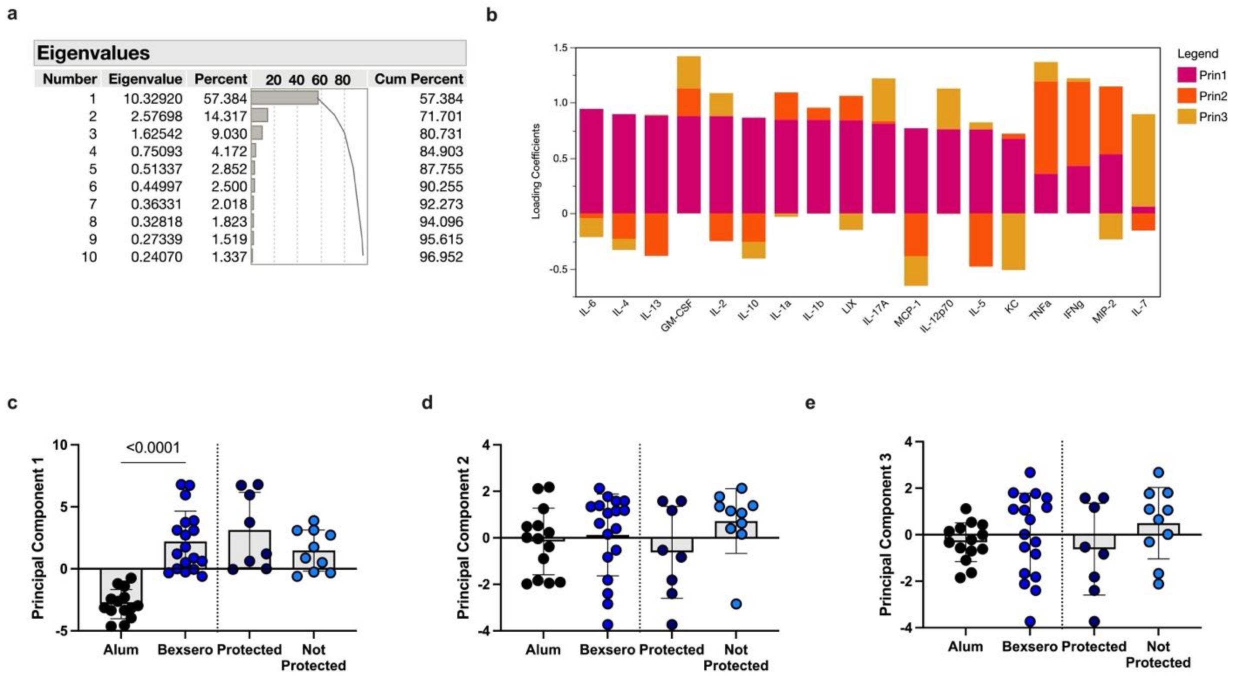


**<u>Supplemental Figure 12.</u> T cell 18-plex Principal Component Analysis results in Bexsero-stimulated spleen samples**. (**a**) The first ten eigenvalues with individual percentages and cumulative percentages. The first three principal components (PC) explain nearly 81% of the variability of the data. (PC1 = 57.4%, PC2 = 14.3%, and PC3 = 9.0%). (**b**) Stack-plot of loading coefficients on the first 3 principal components. The x-axis shows cytokines and chemokines, the y-axis shows the magnitude and direction of the loading coefficients on PC1 (pink), PC2 (orange), and PC3 (yellow) for each variable. Note that individual loadings range between -1 (strongest possible negative correlation) and 1 (strongest possible positive correlation). Scores for PC1 (**c**), PC2 (**d**), and PC3 (**e**) comparing alum (black) and Bexsero (blue) treated animal (left) and protected (dark blue) and not protected (light blue) Bexsero-vaccinated animals (right). Unpaired t-tests compared the groups with p-values shown.

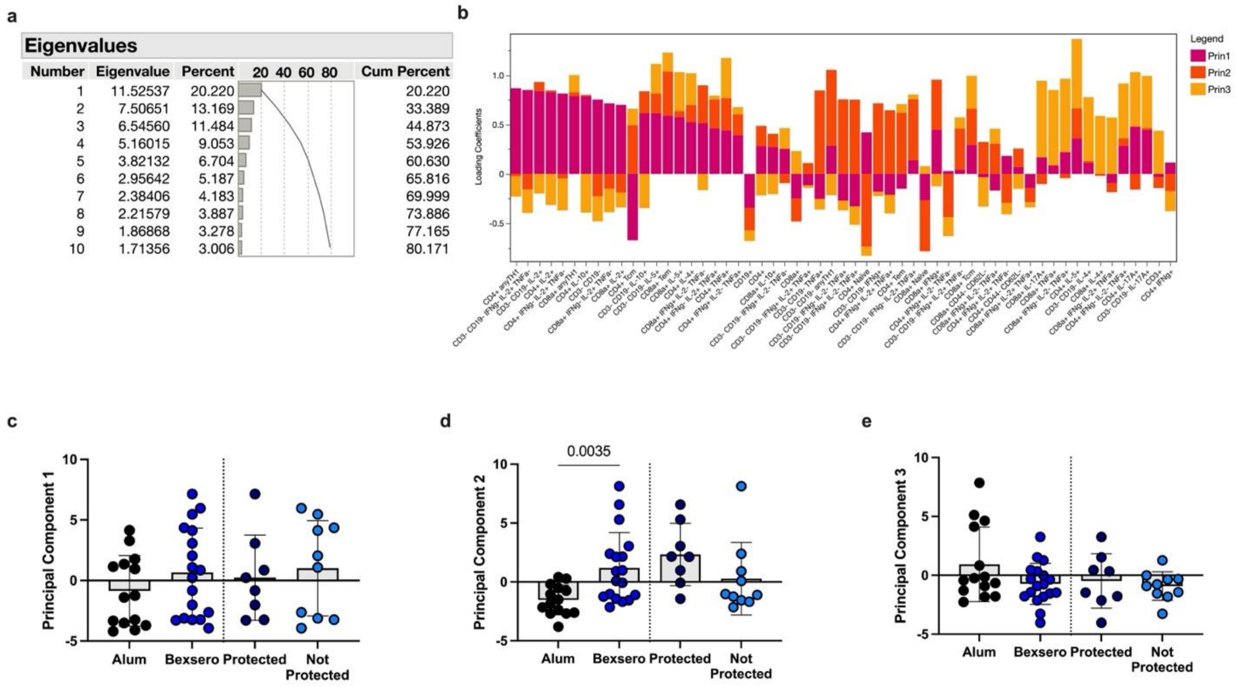


<u>Supplemental Figure 13.</u> **Flow cytometry T cell frequencies Principal Component Analysis results in Bexsero-stimulated splenocytes**. (**a**) The first ten eigenvalues with individual percentages and cumulative percentages. The first three principal components (PC) explain nearly 45% of the variability of the data. (PC1 = 20.22%, PC2 = 13.169%, and PC3 = 11.484%). (**b**) Stack-plot of loading coefficients on the first 3 principal components. The x-axis shows cell subsets as well as cytokine-producing cells, the y-axis shows the magnitude and direction of the loading coefficients on PC1 (pink), PC2 (orange), and PC3 (yellow) for each variable. Note that individual loadings range between -1 (strongest possible negative correlation) and 1 (strongest possible positive correlation). Scores for PC1 (**c**), PC2 (**d**), and PC3 (**e**) comparing alum (black) and Bexsero (blue) treated animal (left) and protected (dark blue) and not protected (light blue) Bexsero-vaccinated animals (right). Unpaired t-tests compared the groups with p-values shown.

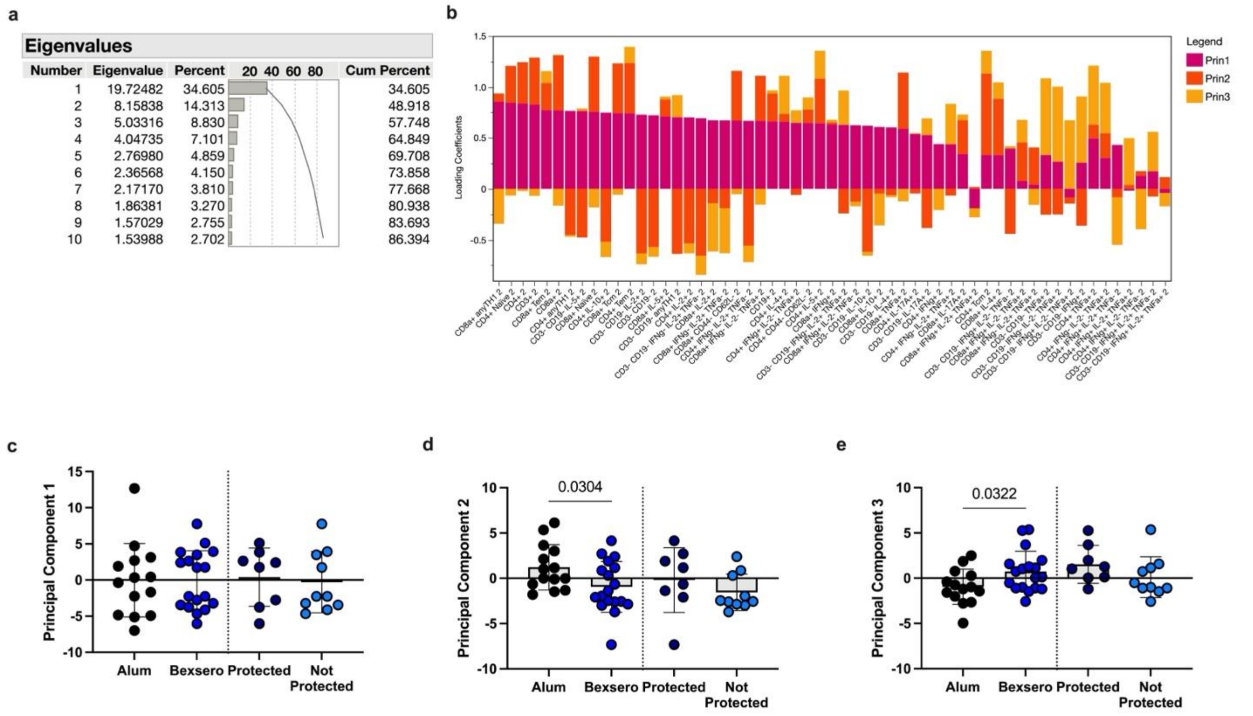


| Number | Eigenvalue | Percent | Cum Percent |
|---|---|---|---|
| 1 | 19.72482 | 34.605 | 34.605 |
| 2 | 8.15838 | 14.313 | 48.918 |
| 3 | 5.03316 | 8.830 | 57.748 |
| 4 | 4.04735 | 7.101 | 64.849 |
| 5 | 2.76980 | 4.859 | 69.708 |
| 6 | 2.36568 | 4.150 | 73.858 |
| 7 | 2.17170 | 3.810 | 77.668 |
| 8 | 1.86381 | 3.270 | 80.938 |
| 9 | 1.57029 | 2.755 | 83.693 |
| 10 | 1.53988 | 2.702 | 86.394 |

<u>Supplemental Figure 14.</u> **Flow cytometry T cell total cells per gram of tissue Principal Component Analysis results in Bexsero-stimulated splenocytes**. (**a**) The first ten eigenvalues with individual percentages and cumulative percentages. The first three principal components (PC) explain nearly 58% of the variability of the data. (PC1 = 34.605%, PC2 = 14.313%, and PC3 = 8.830%). (**b**) Stack-plot of loading coefficients on the first 3 principal components. The x-axis shows cell subsets as well as cytokine-producing cells, the y-axis shows the magnitude and direction of the loading coefficients on PC1 (pink), PC2 (orange), and PC3 (yellow) for each variable. Note that individual loadings range between -1 (strongest possible negative correlation) and 1 (strongest possible positive correlation). Scores for PC1 (**c**), PC2 (**d**), and PC3 (**e**) comparing alum (black) and Bexsero (blue) treated animal (left) and protected (dark blue) and not protected (light blue) Bexsero-vaccinated animals (right). Unpaired t-tests compared the groups with p-values shown.

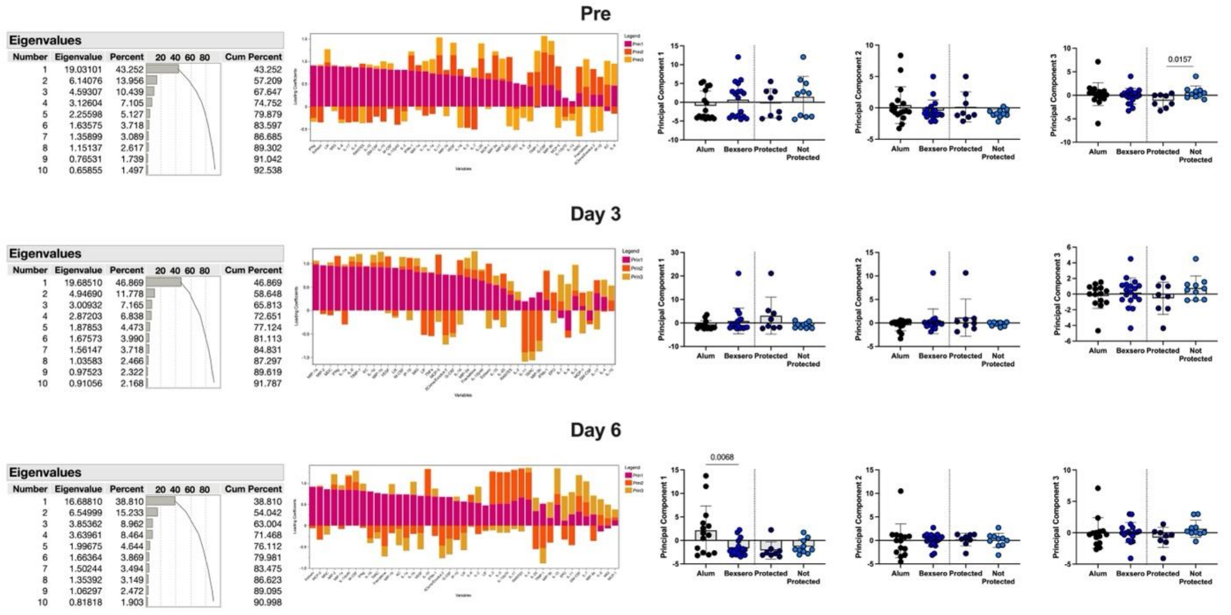


**<u>Supplemental Figure 15.</u> 45-plex Principal Component Analysis results from vaginal lavages**. Left, The first ten eigenvalues with individual percentages and cumulative percentages. Middle, Stack-plot of loading coefficients on the first 3 principal components from each indicated timepoint. The x-axis shows cytokines and chemokines, the y-axis shows the magnitude and direction of the loading coefficients on PC1 (pink), PC2 (orange), and PC3 (yellow) for each variable. Note that individual loadings range between -1 (strongest possible negative correlation) and 1 (strongest possible positive correlation). Right, scores for PC1, PC2, and PC3 comparing alum (black) and Bexsero (blue) treated animal (left) and protected (dark blue) and not protected (light blue) Bexsero-vaccinated animals (right). Unpaired t-tests compared the groups with p-values shown. Principal component analyses from each timepoint (prior to infect but post-vaccination, Pre, Top; day 3 post-infection, Middle; day 6 post-infection, Bottom) were performed independent of one another.
