## supplemental tables for "Meningococcal vaccine Bexsero elicits a robust cellular immune response that targets but is not consistently protective against *Neisseria gonorrhoeae* during murine vaginal infection"

**Supplemental Table 1. Animal information, Bexsero vaccination, gonococcal challenge dose and outcome data**

| Species | Toronto Animal ID | Gender | Vaccine Group | Vaccine Cohort | Infection Cohort | Ngo Infection Dose (CFU) | CFU Recovery | | | | | | | | | | | |
|---|---|---|---|---|---|---|---|---|---|---|---|---|---|---|---|---|---|---|
| | | | | | | | Day 1 | Day 2 | Day 3 | Day 4 | Day 5 | Day 6 | Day 7 | Day 8 | Day 9 | Day 10 | Cumulative CFU | Protection Status |
| BALB/c | 72942 | Female | Alum | 2 | 7 | 1.28 × 10^7 | 2500 | 112 | 37 | 0 | 225 | 2500 | NA | NA | NA | NA | 5374 | NA |
| BALB/c | 72945 | Female | Alum | 2 | 7 | 1.28 × 10^7 | 0 | 112 | 0 | 75 | 8000 | 35000 | NA | NA | NA | NA | 43187 | NA |
| BALB/c | 72946 | Female | Alum | 2 | 7 | 1.28 × 10^7 | 27500 | 1000 | 9000 | 500 | 3750 | 450000 | NA | NA | NA | NA | 491750 | NA |
| BALB/c | 72809 | Female | Bexsero | 2 | 7 | 1.28 × 10^7 | 145000 | 1500 | 150 | 0 | 2500 | 15000 | NA | NA | NA | NA | 164150 | Unprotected |
| BALB/c | 72811 | Female | Bexsero | 2 | 7 | 1.28 × 10^7 | 7000 | 18500 | 375 | 75 | 450 | 40000 | NA | NA | NA | NA | 66400 | Unprotected |
| BALB/c | 72979 | Female | Bexsero | 2 | 7 | 1.28 × 10^7 | 15500 | 0 | 0 | 0 | 0 | 0 | NA | NA | NA | NA | 15500 | Protected |
| BALB/c | 72998 | Female | Bexsero | 2 | 7 | 1.28 × 10^7 | 20000 | 1500 | 0 | 0 | 0 | 0 | NA | NA | NA | NA | 21500 | Protected |
| BALB/c | 72973 | Female | Bexsero | 2 | 7 | 1.28 × 10^7 | 40500 | 4500 | 412 | 112 | 0 | 0 | NA | NA | NA | NA | 45524 | Unprotected |
| BALB/c | 72975 | Female | Bexsero | 2 | 7 | 1.28 × 10^7 | 175000 | 30000 | 0 | 1125 | 2100 | 75000 | NA | NA | NA | NA | 283225 | Unprotected |
| BALB/c | 72943 | Female | Alum | 2 | 8 | 1.33 × 10^7 | 20000 | 150 | 1012 | 12000 | 3000 | 22500 | NA | NA | NA | NA | 58662 | NA |
| BALB/c | 72976 | Female | Bexsero | 2 | 8 | 1.33 × 10^7 | 150000 | 37 | 1312 | 2887 | 187 | 862 | NA | NA | NA | NA | 155285 | Unprotected |
| BALB/c | 72978 | Female | Bexsero | 2 | 8 | 1.33 × 10^7 | 700000 | 2437 | 450 | 750 | 10500 | 225 | NA | NA | NA | NA | 714362 | Unprotected |
| BALB/c | 72947 | Female | Alum | 2 | 9 | 2.50 × 10^6 | 1087 | 500 | 650000 | 225 | 37 | 6500 | NA | NA | NA | NA | 658349 | NA |
| BALB/c | 72950 | Female | Alum | 2 | 9 | 2.50 × 10^6 | 3000 | 37 | 4000 | 11500 | 0 | 37 | NA | NA | NA | NA | 18574 | NA |
| BALB/c | 72825 | Female | Alum | 3 | 10 | 9.00 × 10^7 | 0 | 0 | 90000 | 225 | 75 | 0 | NA | NA | NA | NA | 90300 | NA |
| BALB/c | 72831 | Female | Alum | 3 | 10 | 9.00 × 10^7 | 0 | 0 | 10000 | 14500 | 187 | 150 | NA | NA | NA | NA | 24837 | NA |
| BALB/c | 72835 | Female | Bexsero | 3 | 10 | 9.00 × 10^7 | 0 | 0 | 187 | 0 | 0 | 75 | NA | NA | NA | NA | 262 | Unprotected |
| BALB/c | 72839 | Female | Bexsero | 3 | 10 | 9.00 × 10^7 | 0 | 0 | 0 | 0 | 0 | 0 | NA | NA | NA | NA | 0 | Protected |
| BALB/c | 72848 | Female | Bexsero | 3 | 10 | 9.00 × 10^7 | 0 | 0 | 1600000 | 112 | 0 | 0 | NA | NA | NA | NA | 1600112 | Unprotected |
| BALB/c | 72843 | Female | Bexsero | 3 | 10 | 9.00 × 10^7 | 0 | 0 | 0 | 0 | 0 | 0 | NA | NA | NA | NA | 0 | Protected |
| BALB/c | 72844 | Female | Bexsero | 3 | 10 | 9.00 × 10^7 | 0 | 1425 | 25000 | 0 | 0 | 0 | NA | NA | NA | NA | 26425 | Protected |
| BALB/c | 72845 | Female | Bexsero | 3 | 10 | 9.00 × 10^7 | 0 | 2212 | 100000 | 187 | 412 | 0 | NA | NA | NA | NA | 102811 | Unprotected |
| BALB/c | 72821 | Female | Alum | 3 | 10 | 9.00 × 10^7 | 1000 | 3000 | 1950000 | 0 | 262 | 225 | NA | NA | NA | NA | 1954487 | NA |
| BALB/c | 72823 | Female | Alum | 3 | 10 | 9.00 × 10^7 | 37 | 75 | 4150000 | 200000 | 0 | 0 | NA | NA | NA | NA | 4350112 | NA |
| BALB/c | 72829 | Female | Alum | 3 | 10 | 9.00 × 10^7 | 37 | 0 | 525 | 25000 | 300 | 0 | NA | NA | NA | NA | 25862 | NA |
| BALB/c | 72847 | Female | Bexsero | 3 | 10 | 9.00 × 10^7 | 412 | 3750 | 37 | 12000 | 3000 | 37 | NA | NA | NA | NA | 19236 | Unprotected |
| BALB/c | 72830 | Female | Alum | 3 | 11 | 1.33 × 10^6 | 3750 | 525 | 28000 | 1000 | 1612 | 150 | NA | NA | NA | NA | 35037 | NA |
| BALB/c | 72837 | Female | Bexsero | 3 | 11 | 1.33 × 10^6 | 3750 | 37 | 150 | 0 | 0 | 0 | NA | NA | NA | NA | 3937 | Protected |
| BALB/c | 72840 | Female | Bexsero | 3 | 11 | 1.33 × 10^6 | 225 | 787 | 37 | 0 | 0 | 0 | NA | NA | NA | NA | 1049 | Protected |
| BALB/c | 72819 | Female | Alum | 3 | 12 | 3.00 × 10^6 | 2750000 | 1000 | 150000 | 460000 | 150 | 0 | NA | NA | NA | NA | 3361150 | NA |
| BALB/c | 72828 | Female | Alum | 3 | 12 | 3.00 × 10^6 | 650000 | 37500 | 712 | 525 | 0 | 0 | NA | NA | NA | NA | 688737 | NA |
| BALB/c | 72838 | Female | Bexsero | 3 | 12 | 3.00 × 10^6 | 30000 | 412 | 3000 | 0 | 0 | 0 | NA | NA | NA | NA | 33412 | Protected |

**<u>Supplemental Table 2.</u> Principal component analysis loading table for T cell 18-plex on murine splenocytes**

| Row | Prin1 | Prin2 | Prin3 |
|---|---|---|---|
| IL-6 | 0.943289636 | -0.043821907 | -0.170833675 |
| IL-4 | 0.895954669 | -0.230580793 | -0.099699157 |
| IL-13 | 0.883484331 | -0.384588997 | 0.006755301 |
| GM-CSF | 0.877939408 | 0.249026827 | 0.292384037 |
| IL-2 | 0.877350321 | -0.25045115 | 0.208453501 |
| IL-10 | 0.8644381 | -0.256832273 | -0.152263584 |
| IL-1a | 0.84277425 | 0.248833048 | -0.031517349 |
| IL-1b | 0.842205134 | 0.111773625 | -0.003067613 |
| LIX | 0.839566723 | 0.221072184 | -0.150806098 |
| IL-17A | 0.809319171 | 0.022449869 | 0.386908644 |
| MCP-1 | 0.768708171 | -0.387216444 | -0.267781692 |
| IL-12p70 | 0.758603871 | -0.004511394 | 0.368817918 |
| IL-5 | 0.757137887 | -0.481038992 | 0.064545774 |
| KC | 0.673561995 | 0.045049188 | -0.511988535 |
| MIP-2 | 0.532802609 | 0.612802593 | -0.235756693 |
| IFNg | 0.426819092 | 0.761058312 | 0.03165616 |
| TNFa | 0.354788347 | 0.834911885 | 0.17725357 |
| IL-7 | 0.060329284 | -0.155854709 | 0.835853198 |

**Supplemental Table 3.** **Principal component analysis loading table for T cell flow cytometry frequencies on murine splenocytes**

| Row | Prin1 | Prin2 | Prin3 |
|---|---|---|---|
| CD4+ anyTH1 | 0.8674063 | -0.0217642 | -0.2078815 |
| CD3- CD19- IFNg- IL-2+ TNFa- | 0.84902735 | -0.1563529 | -0.2395747 |
| CD3- CD19- IL-2+ | 0.83579413 | 0.09450065 | -0.1990186 |
| CD4+ IL-2+ | 0.82589688 | 0.01869723 | -0.3143347 |
| CD4+ IFNg- IL-2+ TNFa- | 0.81353456 | -0.0455519 | -0.3240772 |
| CD8a+ anyTH1 | 0.78740457 | 0.03762742 | 0.17563313 |
| CD4+ IL-10+ | 0.78601288 | 0.01416794 | -0.3919617 |
| CD3- CD19- | 0.75332211 | -0.2267836 | -0.253927 |
| CD8a+ IFNg- IL-2+ TNFa- | 0.71382568 | -0.1509275 | -0.2378109 |
| CD8a+ IL-2+ | 0.69896643 | -0.1888485 | -0.152102 |
| CD3- CD19- IL-10+ | 0.61550393 | 0.22092069 | -0.3467713 |
| CD3- CD19- IL-5+ | 0.61237539 | 0.20105342 | 0.29964072 |
| CD8a+ Tem | 0.58838888 | 0.44794344 | 0.19142979 |
| CD8a+ IL-5+ | 0.57433238 | 0.06072769 | 0.3946092 |
| CD4+ IL-4+ | 0.5243067 | 0.17203997 | 0.32248617 |
| CD8a+ IFNg+ IL-2- TNFa- | 0.51334034 | 0.38333646 | -0.1649329 |
| CD4+ IL-17A+ | 0.4785198 | -0.1588456 | 0.5520396 |
| CD4+ IFNg- IL-2+ TNFa+ | 0.46118957 | 0.29193178 | 0.03867878 |
| CD8a+ IFNg+ | 0.44531105 | 0.50852859 | -0.1263573 |
| CD3- CD19- IL-17A+ | 0.4435388 | 0.02077463 | 0.52743594 |
| CD4+ TNFa+ | 0.43900614 | 0.32709301 | 0.40738686 |
| CD4+ Naïve | 0.42028985 | -0.7360654 | -0.0930727 |
| CD4+ IFNg+ IL-2- TNFa+ | 0.39205403 | 0.2095982 | 0.07487042 |
| CD4+ IL-5+ | 0.35745049 | 0.30472352 | 0.70282745 |
| CD8a+ Tcm | 0.29272375 | 0.36633849 | 0.33420347 |
| CD3- CD19- anyTH1 | 0.2844059 | 0.76848127 | -0.2153902 |
| CD4+ IFNg- IL-2- TNFa+ | 0.2826724 | 0.07618526 | 0.55529407 |
| CD4+ | 0.28056273 | 0.20547789 | -0.2188146 |
| CD8a+ IL-10+ | 0.27232433 | 0.13323815 | -0.2062419 |
| CD3- CD19- IFNg+ IL-2- TNFa- | 0.25326992 | -0.0946583 | 0.21040854 |
| CD8a+ IFNg- IL-2- TNFa+ | 0.22008162 | -0.0434914 | 0.74318108 |
| CD4+ IFNg+ IL-2+ TNFa- | 0.18222278 | -0.2941444 | -0.1156949 |
| CD8a+ IL-17A+ | 0.16814362 | -0.1038942 | 0.77474825 |
| CD3- CD19- IFNg- IL-2+ TNFa+ | 0.13781889 | 0.61683002 | 0.04923669 |
| CD4+ IFNg+ | 0.11485059 | -0.1738695 | -0.2049546 |
| CD3- CD19- IL-4+ | 0.11175535 | 0.01741352 | 0.6478133 |
| CD8a+ TNFa+ | 0.08534647 | -0.0014306 | 0.76353397 |
| CD4+ CD44- CD62L- | 0.06904247 | 0.18686702 | -0.1518701 |
| CD3- CD19- IFNg+ IL-2+ TNFa- | 0.04222587 | 0.4163857 | 0.11403434 |
| CD4+ IFNg+ IL-2- TNFa- | 0.02818082 | -0.4374802 | -0.1934157 |
| CD8a+ IL-4+ | -0.0025381 | -0.0174576 | 0.5869671 |
| CD3+ | -0.0308663 | -0.1123871 | 0.43572575 |
| CD8a+ CD44- CD62L- | -0.0340175 | 0.32294268 | -0.2972997 |
| CD8a+ IFNg- IL-2+ TNFa+ | -0.0919487 | -0.0925868 | 0.57117529 |
| CD3- CD19- IFNg+ IL-2+ TNFa+ | -0.1147213 | 0.11012319 | -0.0269493 |
| CD8a+ IFNg+ IL-2+ TNFa+ | -0.1417582 | -0.1430892 | -0.0548126 |
| CD4+ Tem | -0.1502669 | 0.61869307 | 0.08793132 |
| CD8a+ IFNg+ IL-2- TNFa+ | -0.1679632 | 0.30464032 | 0.15269282 |
| CD3- CD19- IFNg+ | -0.1796954 | 0.71473503 | -0.0455412 |
| CD4+ IFNg+ IL-2+ TNFa+ | -0.2121044 | 0.64412088 | -0.1891608 |
| CD8a+ | -0.2478033 | -0.2338905 | 0.23138889 |
| CD3- CD19- TNFa+ | -0.2519909 | 0.8443305 | -0.1083033 |
| CD8a+ Naïve | -0.2664753 | -0.5157222 | 0.07875588 |
| CD3- CD19- IFNg- IL-2- TNFa+ | -0.2707931 | 0.75523422 | -0.098252 |
| CD3- CD19- IFNg+ IL-2- TNFa+ | -0.3286622 | 0.75189789 | -0.1868805 |
| CD19+ | -0.3430653 | -0.2289349 | -0.1077744 |
| CD4+ Tcm | -0.6713977 | 0.49267197 | 0.16664302 |

**<u>Supplemental Table 4.</u> Principal component analysis loading table for T cell flow cytometry cells per gram on murine splenocytes**

| Row | Prin1 | Prin2 | Prin3 |
|---|---|---|---|
| CD8a+ anyTH1 2 | 0.856001332 | 0.079596365 | -0.342859782 |
| CD4+ Naïve 2 | 0.844877634 | 0.362237535 | -0.06638536 |
| CD4+ 2 | 0.836208773 | 0.406647377 | -0.021127684 |
| CD3+ 2 | 0.826190664 | 0.462414174 | -0.068633212 |
| CD8a+ Tem 2 | 0.770706658 | 0.26711334 | 0.116980163 |
| CD8a+ 2 | 0.7694174 | 0.543690031 | -0.165242994 |
| CD4+ anyTH1 2 | 0.76462129 | -0.452033256 | -0.017294482 |
| CD3- CD19- IL-5+ 2 | 0.763070102 | -0.477243017 | 0.025746975 |
| CD8a+ Naïve 2 | 0.755278172 | 0.542619023 | -0.182188107 |
| CD4+ IL-10+ 2 | 0.747976414 | -0.518754385 | -0.150901176 |
| CD8a+ Tcm 2 | 0.741620894 | 0.490560291 | -0.058279809 |
| CD4+ Tem 2 | 0.739913992 | 0.49411281 | 0.158400132 |
| CD3- CD19- IL-2+ 2 | 0.726757523 | -0.633108502 | -0.107295645 |
| CD3- CD19- 2 | 0.720926732 | -0.570603496 | -0.096413753 |
| CD8a+ IL-5+ 2 | 0.709858718 | 0.166852233 | 0.029236166 |
| CD3- CD19- anyTH1 2 | 0.702496131 | -0.638800795 | 0.217188725 |
| CD4+ IL-2+ 2 | 0.700563699 | -0.535660457 | -0.096790281 |
| CD3- CD19- IFNg- IL-2+ TNFa- 2 | 0.692404431 | -0.656614337 | -0.187845676 |
| CD8a+ IL-2+ 2 | 0.673087341 | -0.14073889 | -0.474392039 |
| CD8a+ IFNg- IL-2+ TNFa- 2 | 0.671831614 | -0.190165835 | -0.441595614 |
| CD8a+ CD44- CD62L- 2 | 0.669560584 | 0.488086093 | -0.053365083 |
| CD4+ IFNg- IL-2+ TNFa- 2 | 0.667152618 | -0.557900897 | -0.161123484 |
| CD8a+ IFNg- IL-2- TNFa+ 2 | 0.6660943 | 0.443096871 | -0.155789562 |
| CD19+ 2 | 0.659316958 | 0.275121393 | 0.03191559 |
| CD4+ IL-4+ 2 | 0.657816578 | 0.076613886 | 0.374055244 |
| CD4+ IFNg+ IL-2- TNFa+ 2 | 0.64683113 | -0.060018766 | 0.123905396 |
| CD4+ CD44- CD62L- 2 | 0.645129696 | 0.130993659 | 0.121302934 |
| CD4+ IL-5+ 2 | 0.641794029 | 0.438491935 | 0.275314913 |
| CD8a+ IFNg+ 2 | 0.631456989 | 0.019339029 | 0.027906298 |
| CD3- CD19- IFNg- IL-2+ TNFa+ 2 | 0.627607761 | -0.241082884 | 0.337232934 |
| CD8a+ IFNg+ IL-2- TNFa- 2 | 0.623784741 | -0.126034034 | -0.044982905 |
| CD3- CD19- IL-10+ 2 | 0.618491515 | -0.617670949 | -0.038935293 |
| CD8a+ IL-10+ 2 | 0.607017609 | -0.047133805 | -0.311568932 |
| CD3- CD19- IL-4+ 2 | 0.602855753 | -0.061264222 | -0.020534349 |
| CD8a+ TNFa+ 2 | 0.587983137 | 0.552918035 | -0.123115957 |
| CD4+ IL-17A+ 2 | 0.537601955 | -0.045948163 | 0.009561078 |
| CD3- CD19- IL-17A+ 2 | 0.526821274 | -0.384593589 | 0.164640365 |
| CD4+ TNFa+ 2 | 0.492328208 | 0.13626346 | 0.580669961 |
| CD4+ IFNg+ 2 | 0.438570271 | -0.000506863 | -0.207990344 |
| CD4+ IFNg- IL-2+ TNFa+ 2 | 0.43693774 | -0.066145619 | 0.396444974 |
| CD4+ IFNg+ IL-2+ TNFa- 2 | 0.430845484 | -0.081848991 | -0.468216984 |
| CD3- CD19- IFNg+ IL-2- TNFa- 2 | 0.395083841 | -0.441931865 | 0.023395117 |
| CD8a+ IL-17A+ 2 | 0.339767481 | 0.33387729 | 0.05434836 |
| CD4+ Tcm 2 | 0.331516518 | 0.798534156 | 0.224949347 |
| CD3- CD19- TNFa+ 2 | 0.330985362 | -0.253123136 | 0.75353137 |
| CD8a+ IL-4+ 2 | 0.329985073 | 0.5525292 | 0.162452207 |
| CD4+ IFNg- IL-2- TNFa+ 2 | 0.301580224 | 0.242679598 | 0.496026299 |
| CD3- CD19- IFNg- IL-2- TNFa+ 2 | 0.266001811 | -0.251371194 | 0.736455845 |
| CD3- CD19- IFNg+ 2 | 0.255490712 | -0.362106272 | 0.651399355 |
| CD3- CD19- IFNg+ IL-2+ TNFa- 2 | 0.171643221 | -0.075251674 | 0.387014586 |
| CD4+ IFNg+ IL-2- TNFa- 2 | 0.127151998 | 0.046145398 | -0.396335181 |
| CD8a+ IFNg+ IL-2- TNFa+ 2 | 0.077357083 | 0.376249354 | 0.223504302 |
| CD8a+ IFNg- IL-2+ TNFa+ 2 | 0.039454697 | 0.367675716 | -0.15752943 |
| CD4+ IFNg+ IL-2+ TNFa+ 2 | -0.01716009 | 0.038103019 | 0.459587953 |
| CD3- CD19- IFNg+ IL-2+ TNFa+ 2 | -0.040642931 | 0.115019786 | -0.131354919 |
| CD3- CD19- IFNg+ IL-2- TNFa+ 2 | -0.083519498 | -0.060183482 | 0.672667235 |
| CD8a+ IFNg+ IL-2+ TNFa+ 2 | -0.191106503 | 0.019456918 | -0.088710939 |

<u>Supplemental Table 5.</u> **Principal component analysis loading table for pre-infection vaginal lavage 45-plex**

| Row | Prin1 | Prin2 | Prin3 |
|---|---|---|---|
| IFNg | 0.90946713 | -0.2502848 | -0.0874209 |
| Eotaxin | 0.89572705 | -0.3502643 | 0.03093515 |
| LIX | 0.88958836 | 0.08735584 | 0.29074281 |
| MIG | 0.8841419 | 0.02701388 | -0.0993267 |
| IL-6 | 0.88224633 | -0.2824833 | -0.0806274 |
| IL-17 | 0.87633374 | -0.3528665 | 0.02772044 |
| IL-4 | 0.86601692 | -0.3323215 | -0.0090941 |
| RANTES | 0.86036444 | 0.08638564 | -0.3681361 |
| IL-10 | 0.85958748 | -0.3563974 | 0.06977646 |
| GM-CSF | 0.83387527 | -0.2220909 | 0.06621022 |
| IL-15 | 0.83309531 | -0.3357851 | 0.16662681 |
| M-CSF | 0.81572087 | -0.0512248 | 0.14088938 |
| IL-12p40 | 0.81194578 | -0.3991641 | -0.0056557 |
| IL-2 | 0.8116896 | -0.1045831 | -0.2251445 |
| IFNb-1 | 0.79058842 | 0.33944189 | 0.08882452 |
| MIP-1a | 0.77935898 | 0.13565953 | -0.0782426 |
| IL-1b | 0.77776393 | -0.2017054 | 0.22814581 |
| IL-1a | 0.73448465 | -0.1642332 | 0.31266404 |
| IL-11 | 0.71268199 | 0.42239306 | 0.39344463 |
| MIP-1b | 0.7082589 | 0.10070241 | -0.2110597 |
| VEGF | 0.68209812 | 0.45341628 | 0.2765654 |
| IL-16 | 0.67889719 | 0.14998546 | -0.4688074 |
| IL-3 | 0.66120065 | -0.4658292 | -0.0101923 |
| IL-7 | 0.64011974 | -0.507992 | 0.21259705 |
| IL-20 | 0.61666283 | 0.4469126 | 0.33382967 |
| MCP-1 | 0.60690056 | 0.1336637 | -0.0519188 |
| MIP-3a | 0.59505481 | 0.53669404 | 0.08846214 |
| MIP-2 | 0.57639558 | 0.04006403 | 0.44277739 |
| MDC | 0.55516261 | 0.54061808 | -0.4375876 |
| EPO | 0.51432491 | -0.0512937 | -0.4653358 |
| IL-5 | 0.48283067 | 0.0040343 | -0.251448 |
| MIP-3b | 0.47812529 | 0.66505921 | 0.3085963 |
| G-CSF | 0.47275489 | 0.72163973 | 0.36559175 |
| IL-9 | 0.46232872 | -0.1583802 | 0.48546815 |
| LIF | 0.44635796 | 0.31015984 | -0.3263556 |
| MCP-5 | 0.356887 | 0.53130802 | -0.4710168 |
| TARC | 0.29930198 | 0.57850066 | -0.6572015 |
| 6Ckine/Exodus 2 | 0.26114192 | 0.56906776 | -0.5710094 |
| IP-10 | 0.26081775 | 0.39380202 | -0.5427285 |
| IL-12p70 | 0.19359232 | -0.2246977 | -0.0915913 |
| IL-13 | 0.11881028 | -0.1558545 | -0.1550566 |
| Fractalkine | -0.0012794 | 0.43647907 | 0.61092207 |
| TIMP-1 | -0.0211708 | 0.76598401 | 0.46756 |
| KC | -0.0981059 | 0.48527889 | 0.52524769 |

**Supplemental Table 6. Principal component analysis loading table for day three post-infection vaginal lavage 45-plex**

| Row | Prin1 | Prin2 | Prin3 |
|---|---|---|---|
| MIP-1a | 0.97563113 | 0.02768726 | 0.06169864 |
| MIP-2 | 0.96304099 | -0.0746259 | -0.0051762 |
| MDC | 0.95517627 | -0.1039024 | 0.07247593 |
| IFNg | 0.94279769 | 0.22366468 | -0.0152758 |
| IL-1a | 0.93786007 | -0.2985092 | 0.04660644 |
| IL-6 | 0.93751312 | 0.11246505 | 0.10343022 |
| TIMP-1 | 0.93598331 | 0.09716053 | 0.17268296 |
| KC | 0.93218142 | 0.01659745 | -0.0112575 |
| IL-1b | 0.92759232 | 0.09777748 | 0.1635623 |
| MIP-1b | 0.92401995 | 0.11231615 | 0.2167347 |
| VEGF | 0.91405594 | 0.24227203 | -0.0614426 |
| LIX | 0.91151435 | -0.1106375 | -0.0471488 |
| M-CSF | 0.87543589 | 0.06635323 | 0.20178769 |
| IP-10 | 0.87526712 | 0.24649458 | -0.2081685 |
| MIG | 0.81977968 | 0.12895791 | 0.16724345 |
| LIF | 0.81065416 | -0.5234031 | 0.1139114 |
| TNFa | 0.80419944 | -0.5479006 | 0.01200191 |
| MCP-5 | 0.77353269 | 0.42057781 | -0.2442674 |
| 6Ckine/Exod | 0.76639523 | -0.5277585 | -0.2013148 |
| G-CSF | 0.75776049 | -0.484504 | -0.0858809 |
| IL-16 | 0.75662409 | 0.31698621 | -0.3012281 |
| MIP-3a | 0.71283914 | 0.17412843 | 0.23567062 |
| Fractalkine | 0.67971544 | 0.18945052 | 0.3969123 |
| IL-12p40 | 0.63650055 | 0.20571756 | 0.03359261 |
| Eotaxin | 0.57353214 | 0.424849 | -0.3857188 |
| IL-10 | 0.54238293 | -0.2033687 | 0.2018376 |
| IL-20 | 0.47183343 | 0.31490589 | -0.3611965 |
| MIP-3b | 0.38962114 | -0.6452749 | -0.232185 |
| IL-17 | 0.36959576 | 0.27764969 | -0.4038601 |
| RANTES | 0.33712604 | 0.08889586 | 0.19932444 |
| IL-3 | 0.30679021 | 0.11987566 | 0.54738425 |
| IL-4 | 0.28935628 | -0.0251512 | 0.36545498 |
| TARC | 0.27789046 | -0.8793832 | -0.1865231 |
| IFNb-1 | 0.22660572 | 0.61102455 | 0.02196309 |
| IL-2 | 0.20774049 | 0.0228756 | 0.17153486 |
| IL-11 | 0.19818088 | -0.8963462 | -0.196693 |
| MCP-1 | 0.09598029 | 0.28658113 | -0.4926733 |
| EPO | 0.01699275 | 0.36848718 | -0.2690613 |
| IL-15 | -0.0121196 | 0.21552614 | 0.31853255 |
| IL-7 | -0.1642022 | 0.19702767 | 0.56828014 |
| GM-CSF | -0.2135371 | -0.0802471 | 0.40819437 |
| IL-9 | -0.4306592 | -0.1496929 | 0.56599613 |

**Supplemental Table 7.** **Principal component analysis loading table for day six post-infection vaginal lavage 45-plex**

| Row | Prin1 | Prin2 | Prin3 |
|---|---|---|---|
| Eotaxin | 0.9160767 | -0.054132 | -0.0089283 |
| MCP-5 | 0.91209067 | -0.3304439 | 0.03406657 |
| MDC | 0.85936003 | -0.0102596 | -0.1994393 |
| MIP-2 | 0.85130791 | 0.11958419 | 0.24359205 |
| MIP-1a | 0.84260641 | -0.0775843 | 0.10809156 |
| IL-12p40 | 0.84051117 | 0.31937227 | 0.03472081 |
| M-CSF | 0.83606795 | 0.19392727 | 0.2659232 |
| IFNg | 0.82036362 | -0.0520138 | 0.17302062 |
| IL-16 | 0.80752209 | -0.2985487 | -0.1123689 |
| TARC | 0.76963596 | -0.1566213 | -0.1504516 |
| Fractalkine | 0.74825808 | -0.4855438 | -0.2414762 |
| MIP-1b | 0.738336 | -0.1784202 | 0.24397286 |
| KC | 0.73759133 | -0.0732011 | -0.387594 |
| IL-1a | 0.73667021 | -0.2870747 | -0.3040657 |
| IL-1b | 0.72538836 | -0.1952209 | 0.31150386 |
| VEGF | 0.70366725 | -0.1738801 | -0.3843503 |
| IL-10 | 0.70131962 | 0.62999559 | 0.00857536 |
| IFNb-1 | 0.68554249 | -0.242522 | 0.18934465 |
| 6Ckine/Exodus 2 | 0.67922628 | -0.4699978 | -0.2976415 |
| G-CSF | 0.6763685 | -0.3062354 | -0.3726915 |
| RANTES | 0.66454884 | 0.68389067 | -0.1398879 |
| IL-4 | 0.61546492 | 0.64166841 | 0.10952818 |
| IL-17 | 0.58374828 | 0.7287225 | -0.0834572 |
| IP-10 | 0.57884569 | -0.3371415 | 0.10668472 |
| LIX | 0.56760185 | -0.1672367 | -0.4109865 |
| IL-6 | 0.54864917 | 0.17762406 | -0.034169 |
| IL-2 | 0.53430898 | -0.1991889 | 0.38378043 |
| TIMP-1 | 0.51857649 | -0.5299929 | -0.3612513 |
| IL-5 | 0.51397645 | 0.74834424 | -0.1677359 |
| IL-12p70 | 0.51397645 | 0.74834424 | -0.1677359 |
| IL-3 | 0.50167655 | 0.77624149 | -0.1105882 |
| IL-15 | 0.49243049 | 0.25470871 | 0.60504486 |
| LIF | 0.47625593 | -0.1053837 | -0.026432 |
| IL-20 | 0.34082817 | -0.5539862 | 0.31913464 |
| GM-CSF | 0.33014589 | 0.3698028 | 0.49074013 |
| IL-11 | 0.26895829 | -0.4586531 | 0.52426966 |
| MIP-3a | 0.26592574 | -0.3592316 | 0.44275548 |
| IL-7 | 0.22052524 | 0.33370294 | 0.49008801 |
| MIP-3b | 0.18284574 | -0.401232 | 0.37183613 |
| MCP-1 | 0.12685847 | 0.07054244 | 0.18393301 |
| EPO | 0.11918297 | -0.3288165 | 0.57990118 |
| MIG | -0.0650885 | 0.17463255 | 0.32187099 |
| IL-9 | -0.1384656 | 0.25607228 | 0.37010925 |
